# Super-enhancer impairment is a link between MLL4-inactivated lung tumors and their vulnerability to glycolysis pathway inhibition

**DOI:** 10.1101/507202

**Authors:** Hunain Alam, Ming Tang, Mayinuer Maitituoheti, Shilpa S. Dhar, Samir Amin, Bingnan Gu, Tsai-Yu Chen, Yu-His Lin, Jichao Chen, Florian Muller, Francesco J. DeMayo, Laura Baseler, Kunal Rai, Min Gyu Lee

**Author notes:** Corresponding author Min Gyu Lee. Co-corresponding author Kunal Rai.

## Abstract

Epigenetic modifiers often harbor loss-of-function mutations in lung cancer, but their tumor-suppressive roles are poorly characterized. Here we show that lung-specific loss of the gene encoding the histone methyltransferase MLL4 (alias KMT2D; a COMPASS-like enzyme), which is ranked the most highly inactivated epigenetic modifier in lung cancer, strongly promotes lung adenocarcinoma in mice. *Mll4* loss upregulated tumor-promoting programs, including glycolysis. The pharmacological inhibition of glycolysis preferentially impeded tumorigenic growth of human lung cancer cells bearing *MLL4-*inactivating mutations. *Mll4* loss widely impaired epigenomic signals for super-enhancers and enhancers, including the super-enhancer for the circadian rhythm repressor gene *Per2*, and decreased *Per2* expression. Per2 downregulated several glycolytic pathway genes. These findings uncover a distinct tumor-suppressive epigenetic mechanism in which MLL4 enhances Per2-mediated repression of pro-tumorigenic glycolytic genes via super-enhancer activation to suppress lung adenocarcinoma tumorigenesis and also implicate a glycolysis-targeting strategy as a therapeutic intervention for the treatment of *MLL4-* mutant lung cancer.

## Introduction

Lung cancer is the leading cause of cancer death in men and women. Lung cancer patients suffer from a low overall 5-year survival rate (∼18.1%). Lung cancer is often characterized by gain-of-function mutations and gene amplification of oncogenic kinases (e.g. *K-RAS* and *EGFR*) as well as loss-of-function alterations in tumor suppressor genes (e.g. *TP53* and *LKB1/STK11*) ^1-3^. Much research for lung cancer has focused on kinase signaling pathways, leading to the development of the kinase-targeted therapies, such as EGFR mutant inhibitors and the ALK inhibitors. However, a vast majority of lung cancer patients, who are treated with the kinase-targeted therapies, later undergo tumor relapse and drug resistance ^3, 4^. Recently, the use of immune checkpoint inhibitors (e.g., anti-PD1 and anti-PD-L1) has provided significant survival benefit for lung cancer patients with high PD-L1. However, the overall survival rate of lung cancer patients still remains low ^5, 6^. Moreover, a majority of lung cancer patients do not have a well-defined drug target ^6, 7^. For these reasons, there is a great need for a new mechanistic understanding of lung cancer to be used for a therapeutic approach for lung cancer treatment.

Epigenetic alterations, which represent heritable aberrations in gene expression or cellular phenotype without alterations of DNA sequences, have emerged as a major type of cancer-driving events ^8^. Covalent modifications of DNA and histones play a key role in epigenetic regulation of gene expression and include DNA methylation, histone acetylation, and histone methylation. Interestingly, histone methylation, which can occur at lysine and arginine residues in histones, is linked to either gene activation or silencing, depending on the methylation residues within histones ^9^. Notably, histone methylation modifiers and other epigenetic modifiers are often mutated in lung tumors, including lung adenocarcinoma (LUAD) and lung squamous cell carcinoma (LUSC) responsible for 40%–50% and 25%–30% of lung cancer, respectively ^1, 2, 10-12^. In fact, a substantial portion of such mutations results in the loss-of-function, suggesting tumor-suppressive functions of epigenetic modifiers. However, their tumor-suppressive roles in lung cancer are largely unknown. In the present study, our results show that the histone methyltransferase MLL4 (a COMPASS-like enzyme^13^; also called KMT2D, MLL2, and ALR), which is ranked the most highly inactivated epigenetic modifier in lung tumors, is a key lung tumor suppressor against LUAD. Our findings uncover the molecular mechanism by which the epigenetic modifier MLL4 suppresses LUAD and suggest a mechanism-based therapeutic strategy for treatment of LUAD bearing low MLL4 levels or MLL4’s loss-of-function mutations.

## Results

### Lung-specific loss of *Mll4* strongly promotes *K-Ras^G12D^*-induced LUAD in mice

To search an epigenetic modifier with tumor-suppressive function in lung cancer, we examined which epigenetic modifier highly undergoes genomic alterations related to loss-of-function mutations (i.e., truncations, missense mutations, and insertions/deletions). Our analysis of Pan-lung cancer data (LUAD and LUSC) in the TCGA database showed that the gene encoding MLL4 harbored such genomic alterations in about 14% of lung tumor samples (**Fig. 1a; Supplementary Fig. S1a, b**). Although the mutation frequencies of *MLL3/KMT2C* (16%) were similar to that of MLL4, *MLL4* was ranked higher because truncating mutations were more frequent in *MLL4* than in *KMT2C* (**Fig. 1a**). It should be noted that most of the MLL4 truncations result in loss-of-functions because the catalytic SET domain (5397– 5513 aa) is located at the C-terminus of MLL4. Interestingly, *MLL4* was altered in substantial portions of both LUAD and LUSC samples (**Supplementary Fig. S2a, b**). Because these results indicate that MLL4 is a candidate lung tumor suppressor with high genomic alterations, we chose to characterize the role of MLL4 in lung cancer in subsequent analyses.

**Figure 1:**
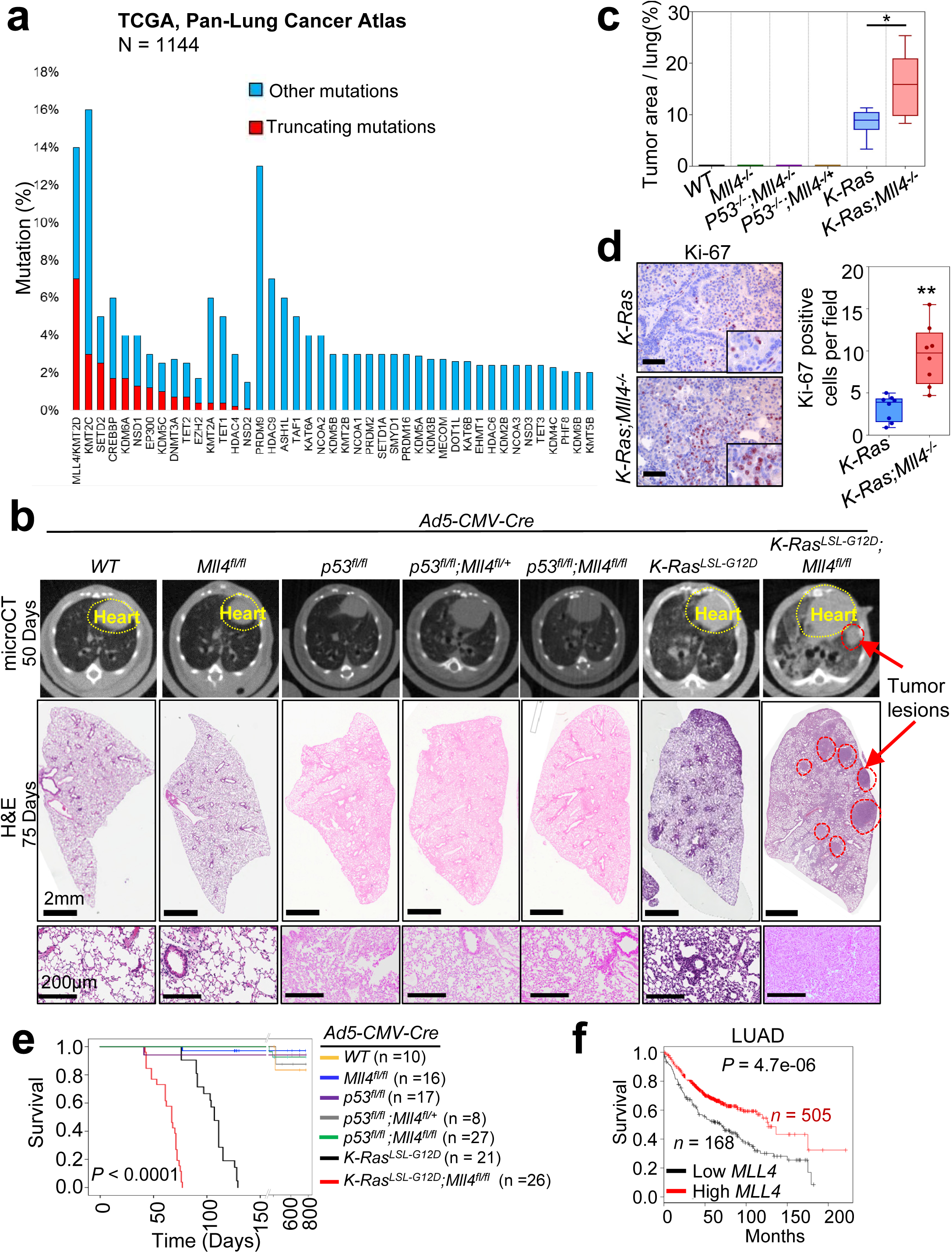
The loss of *Mll4* whose human homologue is the most highly mutated epigenetic modifier in lung cancer accelerates K-Ras-driven LUAD in mice. **(<u>a</u>)** Comparison of alterations of epigenetic modifiers (histone acetyltransferases and deacetylases, histone methyltransferases and demethylases, and DNA modifiers) in TCGA Pan-lung cancer. Other mutations represent missense mutations and inframe mutations. **(<u>b</u>)** Representative micro-CT (top panels) and H&E (middle and bottom panels) images of wild-type (WT), *Mll4*^*fl/fl*^, *p53*^*fl/fl*^, *p53*^*fl/fl*^;*Mll4*^*fl/+*^, *p53*^*fl/fl*^;*Mll4*^*fl/fl*^. *K-Ras*^*LSL-G12D*^, and *K-Ras*^*LSL-G12D*^;*Mll4*^*fl/fl*^ mice. The lungs in mice were infected with Ad5-CMV-Cre viruses. **(<u>c</u>)** Comparison of tumor area (%) per mice in the indicated groups of mice (n = 6). **(<u>d</u>)** IHC analysis for the cell proliferation marker Ki-67 in *K-Ras* and *K-Ras*:*Mll4*^*-/-*^ lung tumors. Ki-67-positive cells in ten random fields of three different tumors of *K-Ras* and *K-Ras*:*Mll4*^*-/-*^ groups were quantified. **(<u>e</u>)** Kaplan-Meier survival analysis of in the indicated groups of mice. **(<u>f</u>)** Kaplan-Meier survival analysis for *MLL4* mRNA levels (probe set, 227527_at) in LUAD patient samples using KM Plotter database (http://kmplot.com/analysis). Lower quartile was used as a cutoff to divide the samples into low and high groups. Scale bars in **b**, 2 mm (middle panels) and 200 µm (bottom panels). *p < 0.05 and **p < 0.01.

*MLL4* alterations frequently co-occur with *p53* and *K-RAS* mutations in lung cancer. For example, about 71% of *MLL4* mutations co-occur with *p53* alterations in lung tumors, and about 22% of *MLL4* mutations coincide with *K-RAS* mutation in LUAD (www.cbioportal.org) (**Supplementary Fig. S3a**). Therefore, we sought to determine whether lung-specific loss of *Mll4* cooperates with *p53* inactivation or K-Ras activation for lung tumorigenesis. We first generated several genetically engineered mouse models (GEMMs), including *p53*^*fl/fl*^;*Mll4*^*fl/fl*^ and *K-Ras*^*LSL-G12D*^;*Mll4*^*fl/fl*^ (**Supplementary Fig. S3b–d**). We then induced Cre-mediated deletion of loxP sites by infecting the lungs of 6–10 weeks old *GEMM* mice with Adeno-Cre viruses via the commonly used intra-tracheal intubation. Lung-specific single loss or co-loss of *Mll4* and *p53* by Adeno-Cre-mediated deletion neither induced any detectable tumor in mouse lungs for up to 16 months post-infection nor changed survival times of mice (**Fig. 1b,c**; **Supplementary Table S1**). This may be partly because *p53* loss alone rarely induces lung tumors in mice ^14-16^. For K-Ras-induced lung tumorigenesis, a *K-Ras* ^*LSL-*G12D^ model (hereinafter referred to as *K-Ras* model) was used because it is a well-established GEMM for LUAD ^17^. *K-Ras* activation was induced by lung-specific deletion of the LSL (loxP-STOP-loxP) cassette via Adeno-Cre virus infection, as previously described ^18^. We monitored potential tumor formation for up to 130 days post-infection at 6–8 week intervals using a micro CT-scan.

Our micro-CT and H&E data showed that lung-specific loss of *Mll4* promoted K-Ras^G12D^-induced lung tumorigenesis (**Fig. 1b, c**). Specifically, our H&E data demonstrated that a higher percentage of the pulmonary parenchyma was effaced by tumors in the *K-Ras;Mll4*^*-/-*^ group of mice than in the *K-Ras* group of mice (**Fig. 1c; Supplementary Fig. S4a, b**). Consistent with tumorigenicity enhanced by *Mll4* loss, immunohistochemisty (IHC) analysis showed that levels of the cell proliferation marker Ki-67 were increased by *Mll4* loss (**Fig. 1d**). Similar to *K-*Ras lung tumors, *K-Ras*:*Mll4*^*-/-*^ lung tumors were positive for the well-known LUAD marker TTF-1 (alias NKX2.1), indicating their LUAD characteristics (**Supplementary Fig. S4c**). In contrast, expression of Keratin 5 (an LUSC marker) was weak in *K-Ras*;*Mll4*^*-/-*^ similar to *K-*Ras tumors (**Supplementary Fig. S4c**). Our survival analysis demonstrated that *Mll4* loss significantly decreased the survival time of mice bearing *K-Ras* pulmonary tumors in the lungs (**Fig. 1e**). Because these results showed that *Mll4* loss accelerated LUAD’s tumorigenicity, we decided to focus on understanding how lung-specific loss of *Mll4* promotes LUAD. In addition, low *MLL4* mRNA levels were associated with poor prognosis in LUAD but not LUSC patients (**Fig. 1f; Supplementary Fig. S4d**).

### Glycolysis program upregulated by *Mll4* loss has human LUAD relevance

To understand how lung-specific loss of *Mll4* promotes LUAD, we isolated tumor lesions from *K-Ras;Mll4*^*-/-*^ and *K-Ras* lungs and examined the effect of *Mll4* loss on expression profiles of *K-Ras* tumors using RNA-seq. Our RNA-seq data confirmed Cre-mediated deletion of loxP in the *Mll4* gene as RNA peaks corresponding to Exon 16–19 of the *Mll4* gene were greatly reduced. (**Supplementary Fig. S5a**). Bioinformatic analyses of RNA-seq data using Gene Set Enrichment Analysis (GSEA) and DAVID tools commonly showed that glycolysis and oxidative phosphorylation pathways were upregulated by *Mll4* loss (**Fig. 2a–d**). Interestingly, many glycolytic genes were also enriched in MYC and mTORC1 pathways. In contrast, these two analyses did not share any common pathway that were significantly downregulated by *Mll4* loss, as GSEA analysis did not indicate any significantly downregulated pathway (**Supplementary Fig. S5b**). In line with our RNA-seq results, analysis of the TCGA lung cancer database showed that glycolysis and oxidative phosphorylation were enriched in human *MLL4*-low/mutant LUAD samples compared with LUAD samples bearing *MLL4*-high levels, indicating that our *K-Ras*:*Mll4*^*-/-*^ tumor model mimics human *MLL4*-low/mutant lung tumors (**Fig. 2e–2g**). Glycolytic genes include *Eno1, Pgk1, Pgam1, Ldha, Gapdh*, and *Cdk1*, whose upregulation by *Mll4* loss was confirmed by quantitative RT-PCR (**Fig. 2h**). In contrast, most genes for oxidative phosphorylation were weakly upregulated by *Mll4* loss (**Fig. 2h**). Therefore, we delved into the regulation of glycolytic genes by MLL4.

**Figure 2:**
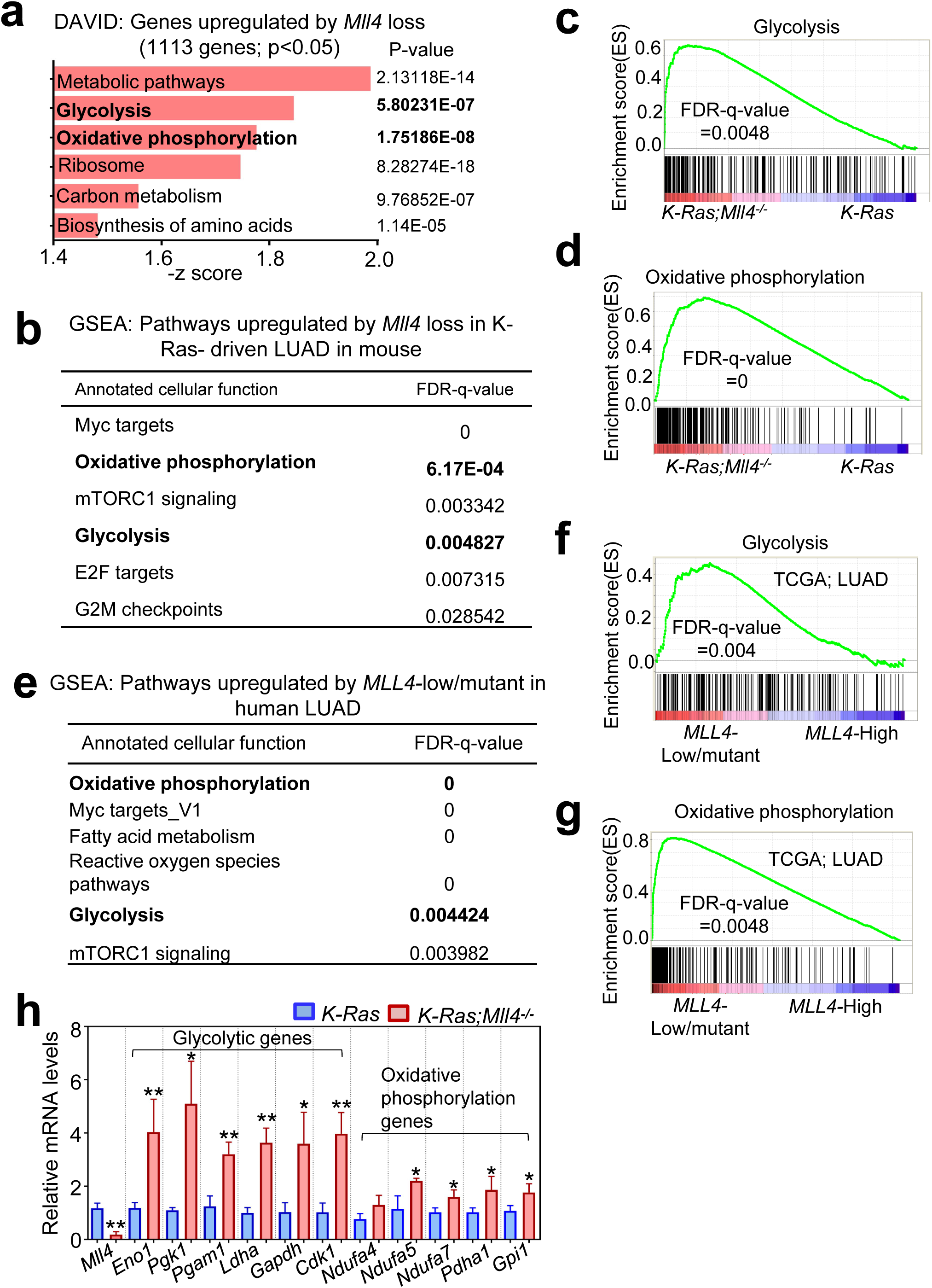
*Mll4* loss upregulates tumor-promoting programs, such as glycolysis. **(<u>a</u>)** Ontology analysis of genes upregulated in *K-Ras*:*Mll4*^*-/-*^ tumors in comparison with *K-Ras* tumors. The functional annotation tool DAVID was used for the analysis. **(<u>b</u>)** GSEA analysis of genes upregulated by *Mll4* loss. **(<u>c and d</u>)** Enrichment plot of glycolysis **(<u>c</u>)** and oxidative phosphorylation **(<u>d</u>)** for genes that were upregulated by *Mll4* loss in mouse LUAD. Each of the black bars represents a gene in the pathway. **(<u>e</u>)** GSEA analysis of genes that were upregulated by in human LUAD samples bearing low *MLL4* expression (n = 20) or *MLL4* inactivation (n = 4) as compared with human LUAD samples (n = 24) bearing high *MLL4* expression. **(<u>f and g</u>)** Enrichment plot of glycolysis **(<u>f</u>)** and oxidative phosphorylation **(<u>g</u>)** in human LUAD samples bearing low *MLL4* expression or *MLL4* inactivation as compared with human LUAD samples bearing high *MLL4* expression. **(<u>h</u>)** Analysis of mRNA levels of glycolytic genes and oxidative phosphorylation-associated genes in *K-Ras* and *K-Ras*:*Mll4*^*-/-*^lung tumors using quantitative RT-PCR. Data are presented as the mean ± SEM (error bars) of at least three independent experiments or biological replicates. *p < 0.05 and **p < 0.01.

It has been shown that upregulation of glycolysis by ENO1^19^, PGK1^20^, PGAM1^21^, LDHA^22^, and GAPDH^23^ promotes tumorigenesis ^24^. CDK1 is a cell cycle gene^25^ and increases glycolysis^26^. Glycolytic enzymes are considered cancer-therapeutic targets ^19, 27^. In line with the above mRNA expression data, IF and IHC analysis showed that ENO1, PGK1, and PGAM1 levels were increased by *Mll4* loss whereas MLL4 levels were decreased (**Fig. 3a–c**). Our analysis of human LUAD database showed that *ENO1, PGK1, PGAM1, LDHA*, and *GAPDH* mRNA levels anti-correlated with *MLL4* mRNA levels in human LUAD samples (**Fig. 3d**). Moreover, high mRNA levels of *ENO1, PGK1, PGAM1, LDHA*, and *GAPDH* were associated with poor survival in LUAD patients (**Fig. 3e**). These results from analysis of human LUAD database indicate human relevance of our findings that *Mll4* loss upregulates expression of glycolytic genes in mouse LUAD.

**Figure 3:**
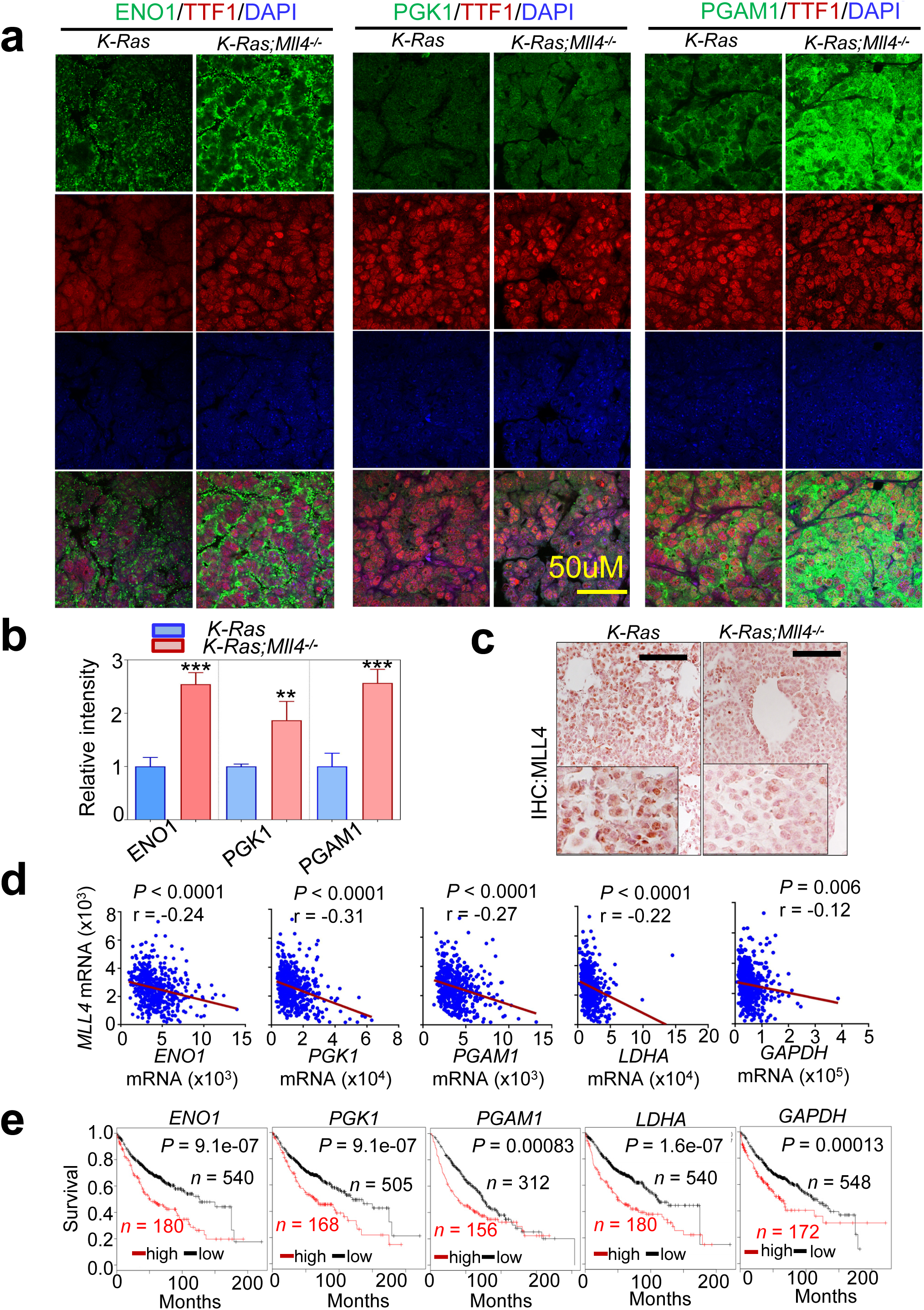
High expression levels of several glycolytic genes anti-correlate with MLL4 levels in LUAD samples and associate with poor prognosis in LUAD patients. **(<u>a and b</u>)** Immunofluorescence (IF) analysis for ENO1, PGK1, and PGAM1 in *K-Ras* and *K-Ras*:*Mll4*^*-/-*^ lung tumor tissues. Representative IF images are shown (**<u>a</u>**), and IF images were quantified (**<u>b</u>**). TTF1 is an LUAD marker. (**<u>c</u>**) Immunohistochemistry (IHC) analysis for MLL4 in *K-Ras* and *K-Ras*:*Mll4*^*-/-*^ lung tumor tissues. (**<u>d</u>**) Anti-correlation of *MLL4* mRNA levels with *ENO1, PGK1, PGAM1, LDHA*, and *GAPDH* mRNA levels in human TCGA LUAD database (n = 517). (**<u>e</u>**) Kaplan-Meier survival analysis for *ENO1, PGK1, PGAM1, LDHA*, and *GAPDH* mRNA levels in human LUAD patients in the KM Plotter database. *ENO1* probe set, 201231_at; *PGK1* probe set, 227068_at; *PGAM1* probe set, 200886_at; *LDHA* probe set, 200650_s_at; *GAPDH* probe set, 212581_at; Scale bar, 50 μm. **p < 0.01 and ***p<0.001

### *Mll4* loss globally reduces super-enhancer and enhancer signals in mouse LUAD

We and others have shown that MLL4 can catalyze mono-, di, and trimethylation at H3K4 ^28-32^. Monomethyl H3K4 (H3K4me1), together with acetyl H3K27 (H3K27ac), marks enhancers, which spatiotemporally activate gene expression in various locations ^33^. MLL4 is required for enhancer formation regulated by the H3K27 acetyltransferases CBP and p300 ^34, 35^. We and others have also shown that MLL4 interacts with the H3K27me3 demethylase UTX ^28, 36, 37^. For these reasons, we performed ChIP-seq for H3K4me1, H3K27ac, H3K4me3, and H3K27me3 using *K-Ras* tumors and *K-Ras*;*Mll4*^*-/-*^ tumors to analyze the effect of *Mll4* loss on epigenomic landscapes and chromatin states. In the ChIP-seq experiment, we also included H3K79me2 and H3K9me3, because H3K79me2 is a commonly analyzed epigenomic mark for gene transcription and H3K9me3 is widely used to study gene repression^38, 39^. Our comprehensive analysis of combinatorial patterns of the 6 histone modifications using the analysis tool ChromHMM ^40^ showed three prominent transitions from K-Ras tumors to *K-Ras*:*Mll4*^*-/-*^ tumors: 1) Active enhancer/State 2 to weak enhancer/State 3; 2) Weak enhancer/State 3 to very low signal/State 9; and 3) Transcribed enhancer/State 7 to H3K4me1-low/State 5 (**Fig. 4a,b; Supplementary Fig. S6a–d**). Further analysis confirmed that *Mll4* loss did not have any obvious effect on H3K4me3 and H3K27me3 signals (**Supplementary Fig. S7a–f**). In line with these results, Western blot, IHC, and IF analyses showed that global levels of H3K4me1 and H3K27ac but not H3K4me3 were downregulated by *Mll4* loss (**Supplementary Fig. S8a,b**). These results indicate that *Mll4* loss highly and negatively impacts enhancer states.

**Figure 4.**
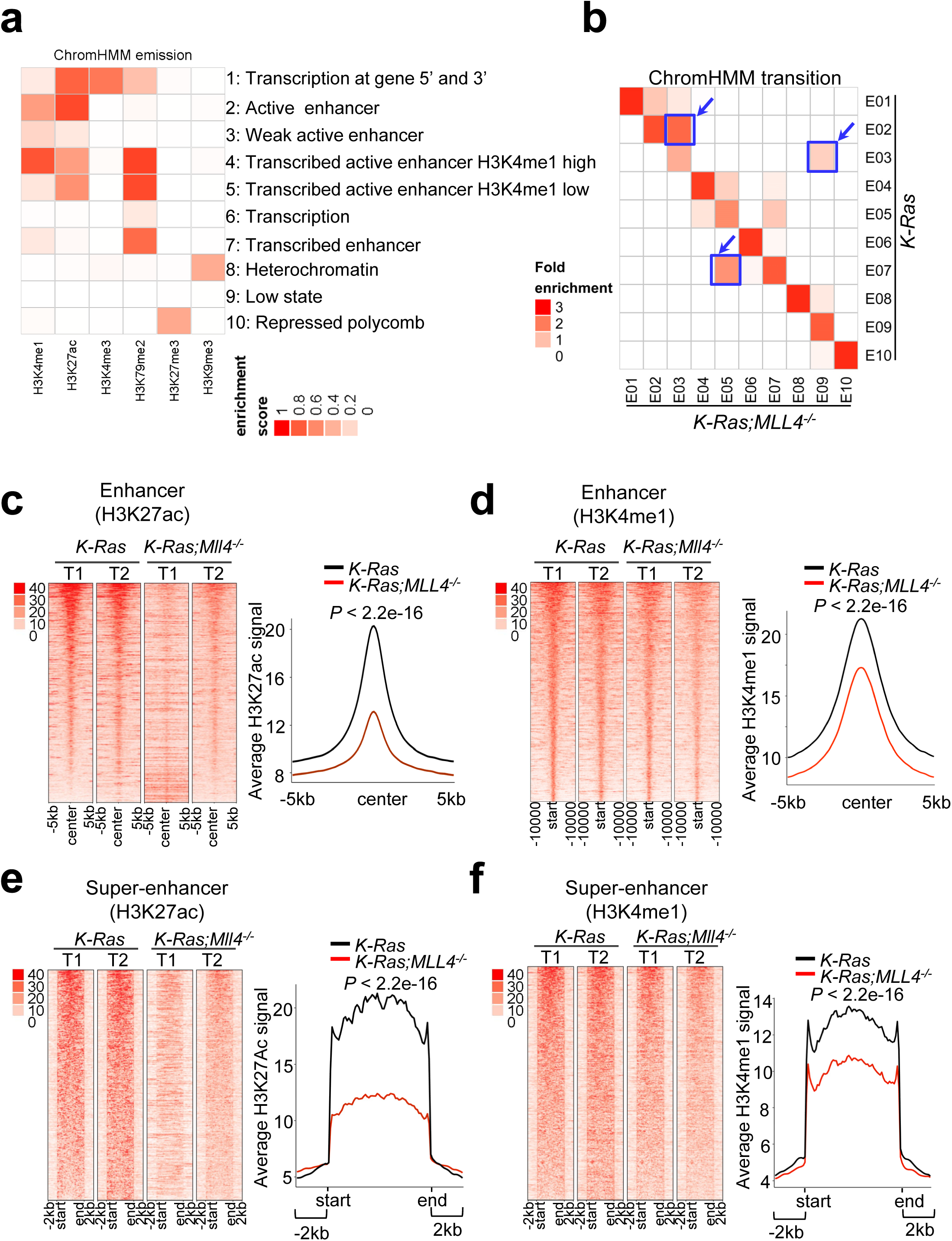
*Mll4* loss reduces epigenomic signals for super-enhancers and to a lesser extent enhancers at the genome-wide level. (**<u>a</u>**) Emission probabilities of the 10-state ChromHMM model calculated from six histone modification profiles in *K-Ras* and *K-Ras*:*Mll4*^*-/-*^ lung tumors. Each row represents one chromatin state. The 10 states predicted using ChromHMM represent various enhancer states (States 2, 3, 4, 5, and 7), promoter state (State 1), transcription (State 1, 6, and 7), polycomb-repressed (state 10), and heterochromatin (State 8). Each column corresponds to a histone modification. The intensity of the color in the scale from 0 (white) to 1 (red) in each cell reflects the frequency of occurrence of each histone mark in the indicated chromatin state. (**<u>b</u>**) Heat map showing the fold enrichment of transitions of chromatin states from *K-Ras* lung tumors to *K-Ras*:*Mll4*^*-/-*^ lung tumors. The color intensities represent the relative fold enrichment. (**<u>c</u>** and **<u>d</u>**) Heat maps (left panels) and average intensity curves (right panels) of ChIP-Seq reads (RPKM) for H3K27ac (**<u>c</u>**) and H3K4me1 (**<u>d</u>**) at typical enhancer regions. Enhancers are shown in a 10kb window centered on the middle of the enhancer) in *K-Ras* and *K-Ras*:*Mll4*^*-/-*^ lung tumors. (**<u>e</u>** and **<u>f</u>**) Heat maps (left panel) and average intensity curves (right panels) of ChIP-Seq reads for H3K27ac (**e**) and H3K4me1 (**<u>f</u>**) at the super-enhancer regions plus their flanking 2kb regions in *K-Ras* and *K-Ras*:*Mll4*^*-/-*^ lung tumors.

It has been documented that enhancers can be categorized into super-enhancers and typical enhancers ^41^. Super-enhancers are characterized by clusters of enhancers that are much larger than typical enhancers with a median size of 0.7–1.3 kb and highly activate gene expression ^41, 42^. Average super-enhancer signals were broader than average typical enhancer signals (**Supplementary Fig. S9a,b**). Interestingly, *Mll4* loss strongly decreased average H3K4me1 and H3K27ac levels in super-enhancers and to a lesser extent typical enhancers (**Fig. 4c–f; Supplementary Fig. S9c,d**). Expression levels of lung-enriched, super-enhancer-associated genes were downregulated (**Supplementary Fig. S9e,f**).

### The pharmacological inhibition of glycolysis impedes the proliferation and tumorigenic growth of LUAD cells with *MLL4*-inactivating mutations

As described above, *Mll4* loss upregulated glycolysis program but decreased H3K27ac levels. This promoted us to test whether the inhibition of glycolytic pathways or H3K27 deacetylation may impede tumorigenic ability of lung tumors bearing *MLL4* loss. For glycolysis inhibition, we used 2-deoxy-D-glucose (2-DG, a hexokinase inhibitor and a glucose analog, and clinical trial phase I/II) ^43, 44^, POMHEX (a new enolase inhibitor) ^a^, koningic acid (KA, a selective inhibitor of GAPDH)^45^, Lonidamine (a hexokinase and mitochondrial respiration inhibitor and clinical trial phase II) ^46^, and Dinaciclib (an inhibitor of CDK1 and CDK2/5/9 and clinical trial phase III) ^47, 48^. To increase histone acetylation via HDAC inhibition, we used AR-42 (Clinical trial phase I) and FDA-approved SAHA/Vorinostat ^49, 50^. We compared the inhibitory effects of these inhibitors on cell-proliferative abilities between human LUAD cell lines with wild type *MLL4* (A549, H1792, H1437, H23, and H358) and those bearing *MLL4’*s truncating mutations (i.e., H1568 with nonsense at E758, DV-90 with the truncating mutation P2118fs, and CORL105 with the truncating mutation R2188fs) (**Supplementary Table S2**). In the *MLL4*-mutant cell lines compared with the *MLL4*-normal cell lines, there were some increased trends of glucose uptake and lactate excretion (**Supplementary Fig. S10a,b**). Our inhibitor experiments showed that 2-DG, POMHEX, KA, and to a lesser extent SAHA and AR-42 inhibited the proliferation of the *MLL4*-mutant cell lines more than the *MLL4*-normal cell lines (**Fig. 5a–f**). In contrast, Lonidamine and Dinaciclib did not exert any selective inhibition against the proliferation of the *MLL4*-mutant cell lines (**Supplementary Fig. S10c, d**).

**Figure 5.**
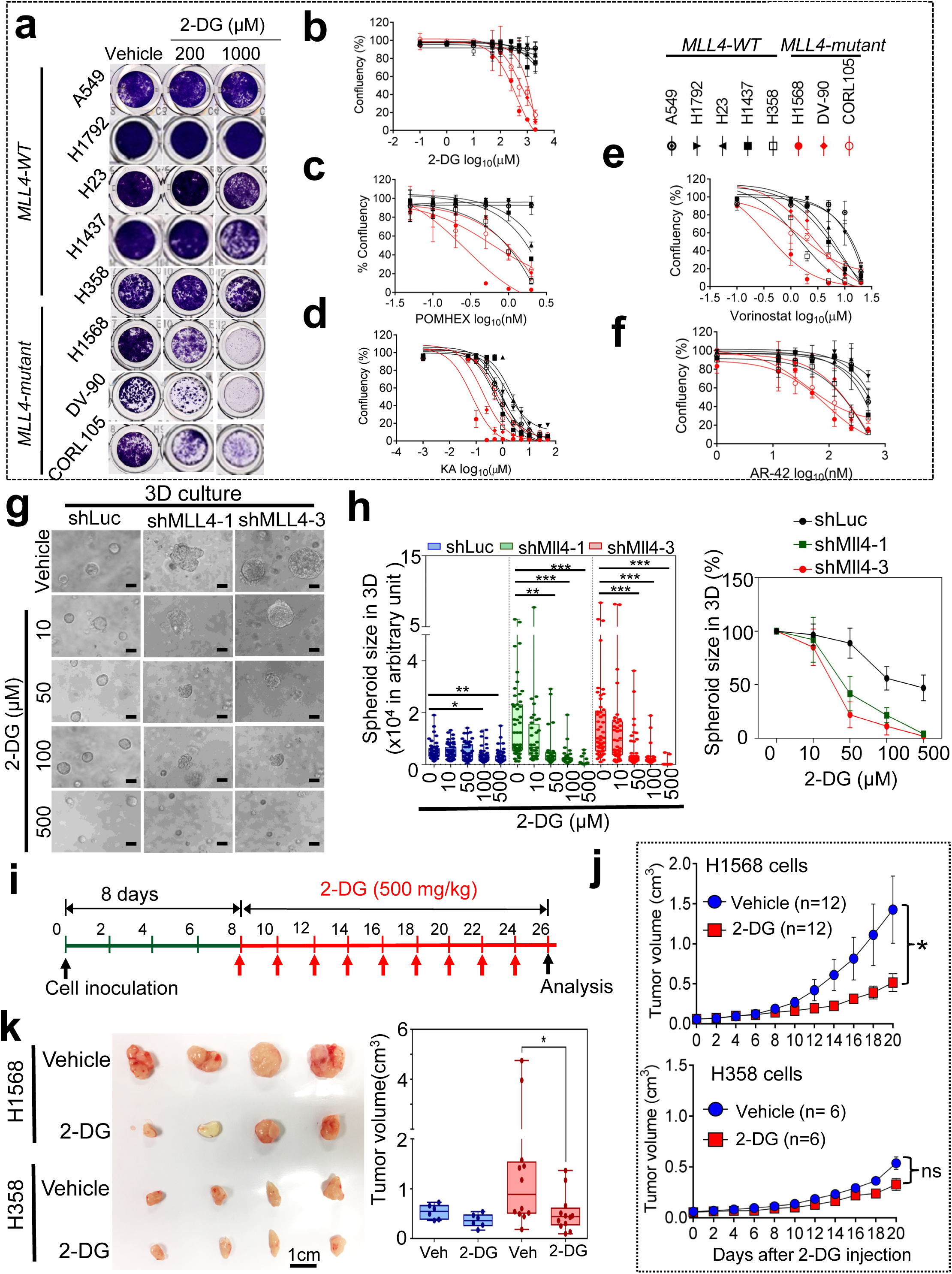
Inhibition of glycolysis suppresses tumorigenic growth of *MLL4*-mutant LUAD cells. (**<u>a</u>**–**<u>f</u>**) Growth inhibition curves of LUAD cell lines bearing wild type (WT) *MLL4* (A549, H1792, H23, H1437, and H358) and those bearing *MLL4* truncation mutants (H1568, DV-90, and CORL105). Cells were treated with various concentrations of 2-DG (**<u>a</u>** and **<u>b</u>**), POMHEX (**<u>c</u>**), KA (**<u>d</u>**), Vorinostat (**<u>e</u>**), and AR-42 (**<u>f</u>**). (**<u>g</u>** & **<u>h</u>**) The effect of 2-DG on spheroid growth of shLuciferase (shLuc)-infected cells and MLL4-depleted LKR-10 cells in a 3D culture. Representative images are shown (**<u>g</u>**). Spheroid sizes were measured (**h**, left panel), and relative sizes of spheroids were plotted (**h**, right panel). (**<u>i</u>**–**<u>k</u>**) The effect of tumorigenic growth of H1568 cells bearing an *MLL4* truncation mutation and of H358 cells bearing wild type MLL4 in a mouse subcutaneous xenograft model. Shown are the treatment schedule of mice with 2-DG (**<u>i</u>**). The sizes of xenograft tumors from H1568 and H358 after the treatment with 2-DG or vehicle control were measured (**<u>j</u>**). Tumors were dissected from the mice after the treatment with 2-DG (500mg/Kg) or vehicle control for 20 days (**<u>k</u>**). Data are presented as the mean ± SEM (error bars) of at least three independent experiments or biological replicates. *p < 0.05, **p < 0.01, and ***p<0.001.

Because 2-DG selectively inhibited the *MLL4*-mutant LUAD cell lines over several *MLL4*-normal cell lines and is currently under clinical trials, we further tested the effect of 2-DG on the growth of the lung cancer cell line LKR10 that is derived from mouse *K-Ras* tumors. A 3D culture system was used, because 1) it better mimics an *in vivo* situation than 2D culture system does, 2) MLL4 knockdown increased the sizes of the spheroids formed from LKR10 cells in the 3D culture (**Supplementary Fig. S10e**), and 3) MLL4 knockdown decreased cell proliferation in the 2D culture system (data not shown). Similar to its inhibitory effect on proliferation of human LUAD cells, 2-DG inhibited more spheroid growth of MLL4-depleted LKR-10 cells than that of control (shLuc)-treated LKR-10 cells in the 3D culture system (**Fig. 5g, h**). In addition, we compared the inhibitory effect of 2-DG on tumorigenic growth between H1568 cells harboring a *MLL4* truncation and H358 cells harboring wild type *MLL4* in a mouse xenograft model. Notably, 2-DG (500 mg/kg per day) preferentially inhibited tumorigenicity of H1568 cells compared with H358 cells (**Fig. 5i–k**).

### MLL4 is required for the activity of the *Per2* super-enhancer, and Per2 represses glycolytic genes

Because MLL4 is a transcriptional co-activator, *Mll4* loss may not directly upregulate tumor-promoting glycolysis program. Thus, we hypothesized that glycolysis program upregulated by *Mll4* loss may be repressed by a tumor-suppressive, transcription-repressive regulator encoded by an MLL4-activated gene (i.e., a gene downregulated by *Mll4* loss). To identify such tumor-suppressive, transcription-repressive regulator, we first searched the overlapping genes between genes downregulated by *Mll4* loss (n = 522) and genes associated with significant decreases in H3K27ac levels (n = 3751), because *Mll4* loss strongly downregulated the enhancer mark H3K27ac to decrease gene expression (**Fig. 6a** and **Supplementary Fig. S11a**). To further reduce the list for such regulator, we then incorporated human lung cancer aspects into our analysis. Specifically, we examined which *Mll4-*loss-downregulated genes correlate with *MLL4* mRNA levels in more than 0.3 of the correlation coefficient value (r) in human LUAD samples in the TCGA database. This analysis resulted in 14 overlapping genes (**Supplementary Fig. S11b,c**). Of these 14 genes, expression levels of five genes were lower in LUAD tumors than in adjacent normal tissues, and their low levels correlated with poor prognosis in lung cancer patients (**Fig. 6b–d; Supplementary Fig. S12a–e**). Interestingly, these genes were occupied by wide H3K27ac and H3K4me1 signals that were decreased by *Mll4* loss (**Fig. 6e; Supplementary Fig. S13a–d**).

**Figure 6.**
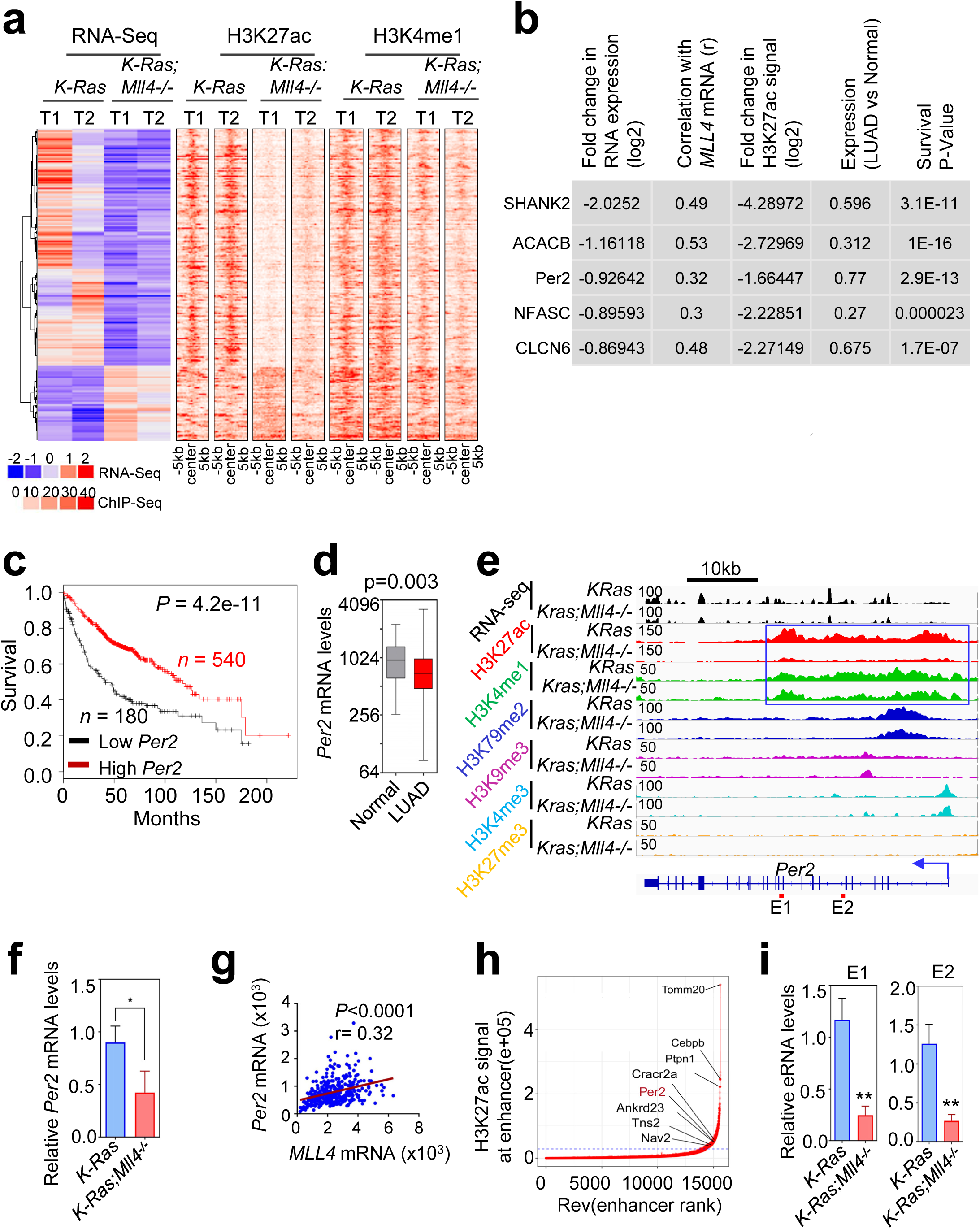
MLL4 positively regulates *Per2* expression. (**<u>a</u>**) Heat maps showing differentially expressed genes and their H3K27ac and H3K4me1 signals in *K-Ras* and *K-Ras*:*Mll4*^*-/-*^ lung tumors. (**<u>b</u>**) The top five hits on the basis of 5 different parameters indicated. (**<u>c</u>**) The Kaplan-Meier survival analysis showing the correlation of low *Per2* mRNA levels with poor survival in lung cancer patients. The auto cutoff was used to divide samples into low and high groups in the KM Plotter database (http://kmplot.com/analysis). Lower quartile was used as a cutoff to divide the samples into low and high groups. *PER2* probe set, 205251_at. (**<u>d</u>**) Box plots showing downregulation of *PER2* mRNA levels in LUAD samples (*n* = 517) compared with their adjacent normal samples (*n* = 54) in TCGA dataset. (**<u>e</u>**) Genome browser view of normalized signals of six chromatin marks (H3K27ac, H3K4me1, H3K79me2, H3K9me3, H3K4me3, and H3K27me3) at the *Per2* locus in *K-Ras* and *K-Ras*:*Mll4*^*-/-*^ lung tumors. All the tracks were average of two biological replicates that were normalized to their inputs. The *Per2*-associated super-enhancer is indicated using a blue line box. (**<u>f</u>**) Analysis of relative *Per2* mRNA levels in *K-Ras* and *K-Ras*:*Mll4*^*-/-*^ lung tumors using quantitative RT-PCR. (**<u>g</u>**) A scatter plot showing a correlation between *MLL4* and *PER2* mRNA levels in TCGA LUAD dataset. (**<u>h</u>**) A plot indicating super-enhancers identified on the basis of H3K27ac signals. (**<u>i</u>**) Analysis of eRNA levels for two different regions (E1 and E2) of the *Per2* super-enhancer using quantitative RT-PCR. Data are presented as the mean ± SEM (error bars) of at least three independent experiments or biological replicates. *p < 0.05 and **p < 0.01.

Of the five genes, the circadian transcriptional repressor gene *Per2* was particularly interesting by the following reasons: 1) *Per2* loss in mouse lung promotes lung tumorigenesis ^51^; 2) *Per2* loss increased glucose metabolism, implicating a role for Per2 in regulating glycolysis ^51^; and 3) disruption of circadian rhythm increases tumorigenicity ^52, 53^. As mentioned above, *Mll4* loss decreased *Per2* expression levels (**Fig. 6f**) and *Per2* mRNA levels significantly correlated with *MLL4* mRNA levels (**Fig. 6g**). Interestingly, *Per2* was occupied by a large cluster of H3K4me1 and H3K27ac peaks that are indicative of a super-enhancer (**Fig. 6e,h**). In enhancer regions, RNAs called enhancer RNAs (eRNAs) are bidirectionally transcribed by RNA Polymerase II ^54^. Enhancer activities can be assessed by eRNA levels ^54^. Therefore, we measured the effect of *Mll4* loss on eRNA levels in the *Per2*–associated super-enhancer. *Mll4* loss decreased eRNA levels in the super-enhancer region, suggesting that *Mll4* loss reduces the activity of the *Per2* super-enhancer (**Fig. 6i**).

To assess whether Per2 represses glycolytic genes that are upregulated by *Mll4* loss, we used the mouse *K-Ras* LUAD line LKR10. We first examined whether MLL4 knockdown in LKR10 cells recapitulates the effect of *Mll4* loss on glycolytic genes in *K-Ras* tumors. In fact, MLL4 knockdown in LKR10 cells upregulated expression levels of several glycolytic genes (e.g., *Eno1, Pgk1, Pgam1, Ldha, Gapdh*, and *Cdk1*) while downregulating *Per2* expression (**Fig. 7a**). In line with this, MLL4 knockdown increased glucose uptake and lactate excretion (**Fig. 7b,c**). We then examined the effect of Per2 knockdown and *Per2* overexpression on expression of these glycolytic genes in LKR10 cells. Per2 knockdown increased expression levels of *Eno1, Pgk1, Pgam1, Ldha, Gapdh*, and *Cdk1* genes, whereas *Per2* overexpression decreased their expression levels (**Fig. 7d,e**). Per2 expression levels anti-correlated with expression of several glycolytic genes (*ENO1, PGK1, PGAM1, LDHA*, and *CDK1*) in LUAD samples in the TCGA database (**Fig. 7f**). Finally, we examined the effect of ectopic expression of *Per2* on the spheroid growth of LKR10 cells that is increased by MLL4 depletion in the 3D culture system. *Per2* re-expression decreased the spheroid growth of MLL4-depleted LKR10 cells (**Fig. 7g**). These results indicate that tumor-suppressive function of MLL4 is dependent, at least in part, on the upregulation of *Per2* expression by MLL4.

**Figure 7.**
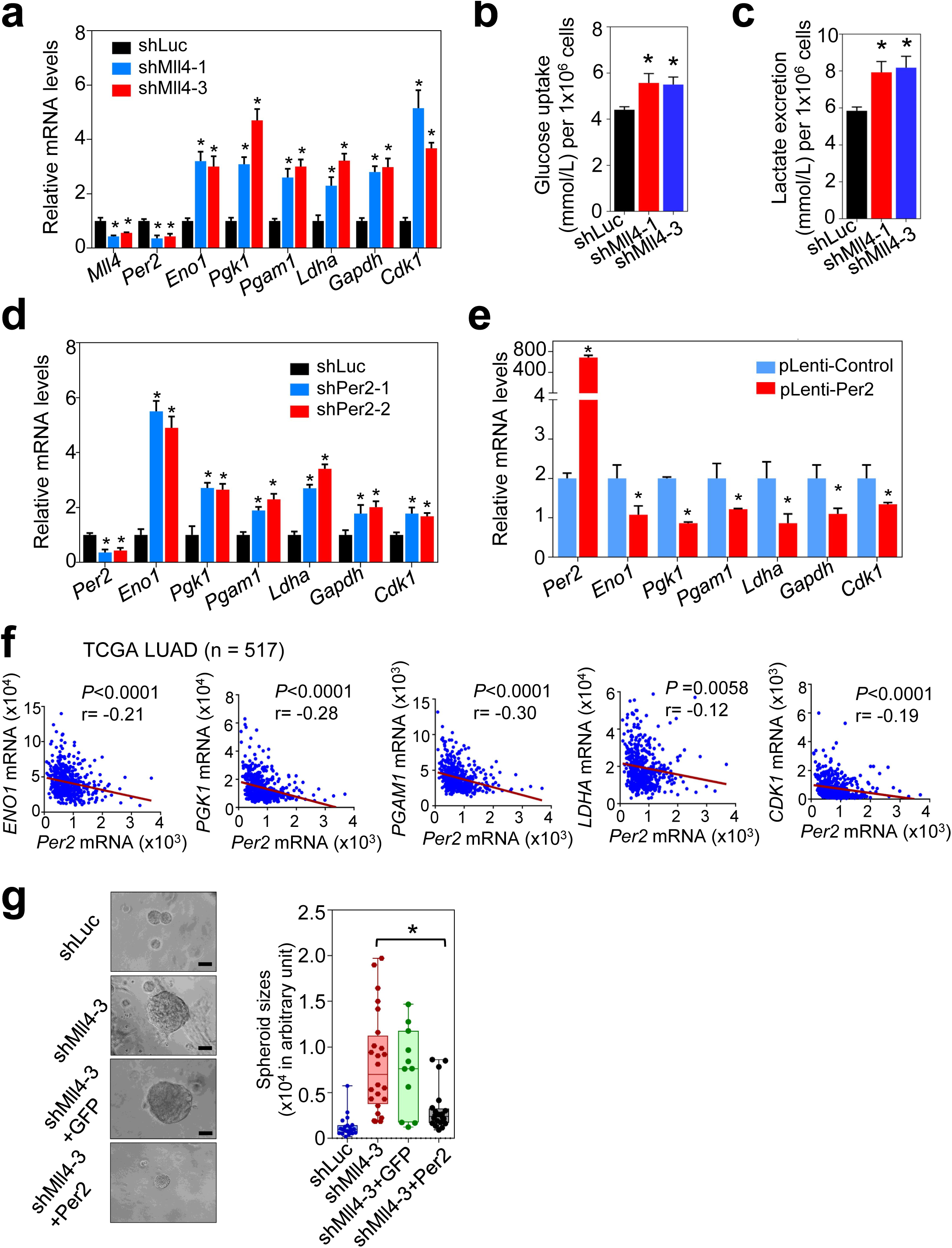
Per2 downregulates glycolytic genes. (**<u>a</u>**) Analysis of the effect of MLL4 knockdown on *Per2, Eno1, Pgk1, Pgam1, Ldha, Gapdh*, and *Cdk1* mRNA levels in mouse LKR-10 cells bearing K-Ras^G12D^ using quantitative RT-PCR. (**<u>b</u>** and **<u>c</u>**) The effect of MLL4 knockdown on glucose uptake (**<u>b</u>**) and lactate excretion (**<u>c</u>**) in LKR10 cells. (**<u>d</u>**) Analysis of the effect of Per*2* knockdown on *Eno1, Pgk1, Pgam1, Ldha, Gapdh*, and *Cdk1* mRNA levels in LKR-10 cells using quantitative RT-PCR. (**<u>e</u>**) Analysis of the effect of ectopic *Per2* expression on *Eno1, Pgk1, Pgam1, Ldha, Gapdh, and Cdk1* mRNA levels in LKR-10 cells using quantitative RT-PCR. (**f**) Scatter plots showing inverse correlations of *PER2* mRNA levels with *ENO1, PGK1, PGAM1, LDHA* or *CDK1 mRNA* levels in human LUAD TCGA dataset (*n* = 517). (**<u>g</u>**) The effect of ectopic expression of Per2 on spheroid sizes of MLL4-depleted LKR-10 cells in 3D-culture. Representative images of spheroid are shown (left panel). The sizes of the spheroids were quantified (right panel). shLuc and pLenti-GFP-infected cells were used as controls. Scale bars represent 200 µm. Data are presented as the mean ± SEM (error bars) of at least three independent experiments or biological replicates. shLuc, shLuciferase *p < 0.05.

## Discussion

It has been reported that MLL4 is required for the formation of acute myeloid leukemia by the MLL-AF9 oncogene ^55^. In contrast, results reported here showed that lung-specific deletion of *Mll4* significantly promoted K-Ras–driven lung tumorigenesis in mice and reduced the survival of mice bearing K-Ras-driven tumors, suggesting that *Mll4* loss cooperates with other oncogenic aberrations to increase LUAD tumorigenicity. The lung tumor-suppressive function of MLL4 is further supported by our following additional results: 1) *Mll4* loss upregulated expression of tumor-promoting glycolytic genes, such as *ENO1, PGK1, LDHA, PGAM1*, and *CDK1*; 2) *Mll4* loss downregulated tumor-suppressive genes, such as *Per2*; and 3) MLL4 depletion increased spheroid sizes of lung cancer cells in the 3D culture system. In line with tumor-suppressive function of MLL4, we have recently shown that brain-specific *Mll4* loss alone induces spontaneous medulloblastoma^56^. Our additional study (the manuscript co-submitted by Rai and colleagues) indicates that MLL4 acts as a tumor suppressor in melanoma tumorigenesis. Other recent studies have demonstrated that genetic ablation of *Mll4* in B cells enhanced B-cell lymphoma genesis, also indicating MLL4’s tumor-suppressive function ^57, 58^. Thus, the anti- or pro-tumorigenicity of MLL4 may be cell-type-dependent, although MLL4 may have a tumor-suppressive function against tumorigenesis in a majority of tissues.

Our results obtained using our *K-Ras*;*Mll4*^*-/-*^ GEMM in the present study indicate that MLL4 acts as an epigenetic LUAD suppressor by positively and highly regulating super-enhancers and to a lesser extent enhancers. Therefore, this GEMM represents a new epigenetic LUAD mouse model. This mouse model would be useful for future additional studies for human LUAD because glycolysis pathway is enriched in both *K-Ras*;*Mll4*^*-/-*^ tumor model and human LUAD tumors with low *MLL4* mRNA levels.

The circadian rhythm transcriptional repressor Per2 plays an important role in tumor suppression ^52, 53^. As our results indicate that the tumor suppressor MLL4 upregulates *Per2* expression to subsequently repress tumor-promoting glycolysis, MLL4-mediated *Per2* activation represents a previously unknown tumor-suppressive mechanism that links between an epigenetic tumor suppressor and a circadian rhythm regulator. Per2 moves into the nucleus during the evening and downregulates gene expression. During the night, Per2 are gradually phosphorylated and targeted for ubiquitination that leads to proteasomal degradation ^59, 60^. Distinct from this type of Per2 regulation, our data suggest a new mode of *Per2*-regulatory mechanism in which MLL4 upregulates *Per2* expression by activating the *Per2* super-enhancer, providing molecular insights into how *Per2* is epigenetically regulated. Super-enhancer formation has been linked to oncogene activation ^61^. However, our results indicate that the *Per2* super-enhancer is associated with tumor-suppressive function in LUAD, consistent with our recent finding that super-enhancer diminution downregulates expression of tumor suppressor genes linked to medulloblastoma genesis ^56^. Our results also define several tumor-promoting glycolytic genes (e.g., *Eno1, Pgk1, Pgam1, Ldha, Gapdh*, and *Cdk1*) as new Per2’s downstream genes. Taken together, our findings reveal a tumor-suppressive mechanism in which MLL4 indirectly represses glycolytic genes by enhancing *Per2* expression via super-enhancer activation and thereby suppresses LUAD.

Increased aerobic glycolysis known as the Warburg effect is a major characteristics of cancer cells and provides cancer cells with proliferation advantage by producing ATP as well as glucose-derived metabolites for biosynthesis of nucleotides, lipids and proteins ^24^. Targeting glycolytic pathways to arrest tumorigenic growth of cancer cells remains an attractive therapeutic intervention. Interestingly, the present study, along with the study co-submitted by Rai and colleagues, showed that *Mll4* loss increased expression of glycolytic genes. In addition, we demonstrated that the glycolysis inhibitor 2-DG impeded tumorigenicity of LUAD cells bearing an *MLL4* mutation in a mouse xenograft model. Because *MLL4* is mutated in other several types of cancer ^62^ and certain glycolysis inhibitor (i.e., 2-DG) has re-entered into clinical trials, our findings may rationalize glycolysis inhibition as an anti-cancer treatment strategy against human *MLL4*-inactivated LUAD and other cancer types bearing MLL4-inactivating mutations.

## Methods

Methods, including experimental procedures and any associated accession numbers and references, are available as supplementary information.

## Acknowledgments

We are thankful to Julien Sage and Tyler Jacks for providing their reagents and also thank Haoqiang Ying, Zhenbo Han, Su Zhang, and Charles Kingsley for their technical assistance. This work was supported by grants to MG Lee from the National Institutes of Health (NIH; R01 CA157919, R01 CA207109, and R01 CA207098) and the Center for Cancer Epigenetics at the UT MD Anderson Cancer Center, by a grant to K Rai from the NIH (CA160578), by a grant to FL Muller from American Cancer Society (RSG-15-145-01-CDD), by a grant to FJ DeMayo from the Intramural Research Program of the National Institute of Environmental Health Sciences (Z1AES103311-10), and by a fellowship to H Alam from the Odyssey program at the UT MD Anderson Cancer Center. The animal imagining and histology work were performed at the Small Animal Imaging Facility and Histopathology Core Lab, respectively, at the UT MD Anderson Cancer Center supported by the NIH National Cancer Institute (P30CA016672).

## Author Contributions

H.A. planned and carried out experiments, analyzed data, prepared figures, and wrote the manuscript. M.T. performed the bioinformatics analysis of RNA-seq and ChIP-seq data, prepared bioinformatics figures, and wrote the manuscript. M.M. and S.S.D. performed ChIP-seq experiments. S.A. contributes to data analysis. T-Y.C. contributed to inhibitor experiments. J.C. contributed to IHC experiments and analysis. Y-H.L., F.M., and F.J.D. provided reagents. L.B. analyzed tumor data. K.R. directed ChIP-seq experiments, guided the bioinformatics study, contributed to experiment design, and wrote the manuscript. M.G.L. conceived the study, designed and oversaw the study, evaluated data, and wrote the manuscript.

## Footnotes

a Lin *et al.*, Eradication of ENO1-deleted Glioblastoma through Collateral Lethality bioRxiv 331538; doi: https://doi.org/10.1101/331538

## References

1. Imielinski, M. et al. Mapping the hallmarks of lung adenocarcinoma with massively parallel sequencing. Cell 150, 1107–1120, doi:10.1016/j.cell.2012.08.029 (2012).

2. Cancer Genome Atlas Research, N. Comprehensive molecular profiling of lung adenocarcinoma. Nature 511, 543–550, doi:10.1038/nature13385 (2014).

3. Herbst, R. S., Heymach, J. V. & Lippman, S. M. Lung cancer. N Engl J Med 359, 1367– 1380, doi:10.1056/NEJMra0802714 (2008).

4. Sharma, S. V., Bell, D. W., Settleman, J. & Haber, D. A. Epidermal growth factor receptor mutations in lung cancer. Nature reviews. Cancer 7, 169–181, doi:10.1038/nrc2088 (2007).

5. Malhotra, J., Jabbour, S. K. & Aisner, J. Current state of immunotherapy for non-small cell lung cancer. Transl Lung Cancer Res 6, 196–211, doi:10.21037/tlcr.2017.03.01 (2017).

6. Herbst, R. S., Morgensztern, D. & Boshoff, C. The biology and management of non-small cell lung cancer. Nature 553, 446–454, doi:10.1038/nature25183 (2018).

7. Chan, B. A. & Hughes, B. G. Targeted therapy for non-small cell lung cancer: current standards and the promise of the future. Transl Lung Cancer Res 4, 36–54, doi:10.3978/j.issn.2218-6751.2014.05.01 (2015).

8. Dawson, M. A. & Kouzarides, T. Cancer epigenetics: from mechanism to therapy. Cell 150, 12–27, doi:S0092-8674(12)00762-3 [pii] 10.1016/j.cell.2012.06.013 (2012).

9. Barski, A. et al. High-resolution profiling of histone methylations in the human genome. Cell 129, 823–837, doi:S0092-8674(07)00600-9 [pii] 10.1016/j.cell.2007.05.009 (2007).

10. Kandoth, C. et al. Mutational landscape and significance across 12 major cancer types. Nature 502, 333–339, doi:10.1038/nature12634 (2013).

11. Cancer Genome Atlas Research, N. Comprehensive genomic characterization of squamous cell lung cancers. Nature 489, 519–525, doi:10.1038/nature11404 (2012).

12. Campbell, J. D. et al. Distinct patterns of somatic genome alterations in lung adenocarcinomas and squamous cell carcinomas. Nat Genet 48, 607–616, doi:10.1038/ng.3564 (2016).

13. Shilatifard, A. The COMPASS family of histone H3K4 methylases: mechanisms of regulation in development and disease pathogenesis. Annu Rev Biochem 81, 65–95, doi:10.1146/annurev-biochem-051710-134100 (2012).

14. Donehower, L. A. et al. Mice deficient for p53 are developmentally normal but susceptible to spontaneous tumours. Nature 356, 215–221, doi:10.1038/356215a0 (1992).

15. Olive, K. P. et al. Mutant p53 gain of function in two mouse models of Li-Fraumeni syndrome. Cell 119, 847–860, doi:10.1016/j.cell.2004.11.004 (2004).

16. Lang, G. A. et al. Gain of function of a p53 hot spot mutation in a mouse model of Li-Fraumeni syndrome. Cell 119, 861–872, doi:10.1016/j.cell.2004.11.006 (2004).

17. DuPage, M., Dooley, A. L. & Jacks, T. Conditional mouse lung cancer models using adenoviral or lentiviral delivery of Cre recombinase. Nat Protoc 4, 1064–1072, doi:10.1038/nprot.2009.95 (2009).

18. Jackson, E. L. et al. Analysis of lung tumor initiation and progression using conditional expression of oncogenic K-ras. Genes Dev 15, 3243–3248, doi:10.1101/gad.943001 (2001).

19. Capello, M., Ferri-Borgogno, S., Cappello, P. & Novelli, F. alpha-Enolase: a promising therapeutic and diagnostic tumor target. FEBS J 278, 1064–1074, doi:10.1111/j.1742-4658.2011.08025.x (2011).

20. Li, X. et al. Mitochondria-Translocated PGK1 Functions as a Protein Kinase to Coordinate Glycolysis and the TCA Cycle in Tumorigenesis. Molecular cell 61, 705–719, doi:10.1016/j.molcel.2016.02.009 (2016).

21. Hitosugi, T. et al. Phosphoglycerate mutase 1 coordinates glycolysis and biosynthesis to promote tumor growth. Cancer Cell 22, 585–600, doi:10.1016/j.ccr.2012.09.020 (2012).

22. Sheng, S. L. et al. Knockdown of lactate dehydrogenase A suppresses tumor growth and metastasis of human hepatocellular carcinoma. FEBS J 279, 3898–3910, doi:10.1111/j.1742-4658.2012.08748.x (2012).

23. Colell, A., Green, D. R. & Ricci, J. E. Novel roles for GAPDH in cell death and carcinogenesis. Cell Death Differ 16, 1573–1581, doi:10.1038/cdd.2009.137 (2009).

24. Gatenby, R. A. & Gillies, R. J. Why do cancers have high aerobic glycolysis? Nature reviews. Cancer 4, 891–899, doi:10.1038/nrc1478 (2004).

25. Malumbres, M. & Barbacid, M. Cell cycle, CDKs and cancer: a changing paradigm. Nature reviews. Cancer 9, 153–166, doi:10.1038/nrc2602 (2009).

26. Wang, X. Q. et al. CDK1-PDK1-PI3K/Akt signaling pathway regulates embryonic and induced pluripotency. Cell Death Differ 24, 38–48, doi:10.1038/cdd.2016.84 (2017).

27. Ganapathy-Kanniappan, S., Kunjithapatham, R. & Geschwind, J. F. Anticancer efficacy of the metabolic blocker 3-bromopyruvate: specific molecular targeting. Anticancer research 33, 13–20 (2013).

28. Lee, M. G. et al. Demethylation of H3K27 regulates polycomb recruitment and H2A ubiquitination. Science 318, 447–450, doi:10.1126/science.1149042 (2007).

29. Dhar, S. S. et al. Trans-tail regulation of MLL4-catalyzed H3K4 methylation by H4R3 symmetric dimethylation is mediated by a tandem PHD of MLL4. Genes Dev 26, 2749– 2762, doi:10.1101/gad.203356.112 (2012).

30. Zhang, Y. et al. Evolving Catalytic Properties of the MLL Family SET Domain. Structure 23, 1921–1933, doi:10.1016/j.str.2015.07.018 (2015).

31. Lee, J. E. et al. H3K4 mono-and di-methyltransferase MLL4 is required for enhancer activation during cell differentiation. Elife 2, e01503, doi:10.7554/eLife.01503 (2013).

32. Herz, H. M. et al. Enhancer-associated H3K4 monomethylation by Trithorax-related, the Drosophila homolog of mammalian Mll3/Mll4. Genes Dev 26, 2604–2620, doi:10.1101/gad.201327.112 (2012).

33. Smith, E. & Shilatifard, A. Enhancer biology and enhanceropathies. Nature structural & molecular biology 21, 210–219, doi:10.1038/nsmb.2784 (2014).

34. Lai, B. et al. MLL3/MLL4 are required for CBP/p300 binding on enhancers and super-enhancer formation in brown adipogenesis. Nucleic acids research 45, 6388–6403, doi:10.1093/nar/gkx234 (2017).

35. Wang, S. P. et al. A UTX-MLL4-p300 Transcriptional Regulatory Network Coordinately Shapes Active Enhancer Landscapes for Eliciting Transcription. Molecular cell 67, 308– 321 e306, doi:10.1016/j.molcel.2017.06.028 (2017).

36. Issaeva, I. et al. Knockdown of ALR (MLL2) reveals ALR target genes and leads to alterations in cell adhesion and growth. Mol Cell Biol 27, 1889–1903, doi:MCB.01506-06 [pii] 10.1128/MCB.01506-06 (2007).

37. Cho, Y. W. et al. PTIP associates with MLL3- and MLL4-containing histone H3 lysine 4 methyltransferase complex. The Journal of biological chemistry 282, 20395–20406 (2007).

38. Nguyen, A. T. & Zhang, Y. The diverse functions of Dot1 and H3K79 methylation. Genes Dev 25, 1345–1358, doi:10.1101/gad.2057811 (2011).

39. Hawkins, R. D. et al. Distinct epigenomic landscapes of pluripotent and lineage-committed human cells. Cell Stem Cell 6, 479–491, doi:10.1016/j.stem.2010.03.018 (2010).

40. Ernst, J. & Kellis, M. ChromHMM: automating chromatin-state discovery and characterization. Nat Methods 9, 215–216, doi:10.1038/nmeth.1906 (2012).

41. Whyte, W. A. et al. Master transcription factors and mediator establish super-enhancers at key cell identity genes. Cell 153, 307–319, doi:10.1016/j.cell.2013.03.035 (2013).

42. Loven, J. et al. Selective inhibition of tumor oncogenes by disruption of super-enhancers. Cell 153, 320–334, doi:10.1016/j.cell.2013.03.036 (2013).

43. Huang, C. C. et al. Glycolytic inhibitor 2-deoxyglucose simultaneously targets cancer and endothelial cells to suppress neuroblastoma growth in mice. Dis Model Mech 8, 1247– 1254, doi:10.1242/dmm.021667 (2015).

44. Pelicano, H., Martin, D. S., Xu, R. H. & Huang, P. Glycolysis inhibition for anticancer treatment. Oncogene 25, 4633–4646, doi:10.1038/sj.onc.1209597 (2006).

45. Liberti, M. V. et al. A Predictive Model for Selective Targeting of the Warburg Effect through GAPDH Inhibition with a Natural Product. Cell Metab 26, 648–659 e648, doi:10.1016/j.cmet.2017.08.017 (2017).

46. Brawer, M. K. Lonidamine: basic science and rationale for treatment of prostatic proliferative disorders. Rev Urol 7 Suppl 7, S21–26 (2005).

47. Parry, D. et al. Dinaciclib (SCH 727965), a novel and potent cyclin-dependent kinase inhibitor. Molecular cancer therapeutics 9, 2344–2353, doi:10.1158/1535-7163.MCT-10-0324 (2010).

48. Santo, L. et al. AT7519, A novel small molecule multi-cyclin-dependent kinase inhibitor, induces apoptosis in multiple myeloma via GSK-3beta activation and RNA polymerase II inhibition. Oncogene 29, 2325–2336, doi:10.1038/onc.2009.510 (2010).

49. Bjornsson, H. T. et al. Histone deacetylase inhibition rescues structural and functional brain deficits in a mouse model of Kabuki syndrome. Science translational medicine 6, 256ra135, doi:10.1126/scitranslmed.3009278 (2014).

50. Jacob, A. et al. Preclinical validation of AR42, a novel histone deacetylase inhibitor, as treatment for vestibular schwannomas. The Laryngoscope 122, 174–189, doi:10.1002/lary.22392 (2012).

51. Papagiannakopoulos, T. et al. Circadian Rhythm Disruption Promotes Lung Tumorigenesis. Cell Metab 24, 324–331, doi:10.1016/j.cmet.2016.07.001 (2016).

52. Filipski, E. et al. Disruption of circadian coordination accelerates malignant growth in mice. Pathologie-biologie 51, 216–219 (2003).

53. Fu, L., Pelicano, H., Liu, J., Huang, P. & Lee, C. The circadian gene Period2 plays an important role in tumor suppression and DNA damage response in vivo. Cell 111, 41–50 (2002).

54. Kim, T. K. & Shiekhattar, R. Architectural and Functional Commonalities between Enhancers and Promoters. Cell 162, 948–959, doi:10.1016/j.cell.2015.08.008 (2015).

55. Santos, M. A. et al. DNA-damage-induced differentiation of leukaemic cells as an anti-cancer barrier. Nature 514, 107–111, doi:10.1038/nature13483 (2014).

56. Dhar, S. S. et al. MLL4 Is Required to Maintain Broad H3K4me3 Peaks and Super-Enhancers at Tumor Suppressor Genes. Molecular cell 70, 825–841 e826, doi:10.1016/j.molcel.2018.04.028 (2018).

57. Ortega-Molina, A. et al. The histone lysine methyltransferase KMT2D sustains a gene expression program that represses B cell lymphoma development. Nat Med, doi:10.1038/nm.3943 (2015).

58. Zhang, J. et al. Disruption of KMT2D perturbs germinal center B cell development and promotes lymphomagenesis. Nat Med, doi:10.1038/nm.3940 (2015).

59. Bass, J. & Takahashi, J. S. Circadian integration of metabolism and energetics. Science 330, 1349–1354, doi:10.1126/science.1195027 (2010).

60. Koike, N. et al. Transcriptional architecture and chromatin landscape of the core circadian clock in mammals. Science 338, 349–354, doi:10.1126/science.1226339 (2012).

61. Sur, I. & Taipale, J. The role of enhancers in cancer. Nature reviews. Cancer 16, 483–493, doi:10.1038/nrc.2016.62 (2016).

62. Suva, M. L., Riggi, N. & Bernstein, B. E. Epigenetic reprogramming in cancer. Science 339, 1567–1570, doi:10.1126/science.1230184 (2013).

