## Supplemental files for "Super-enhancer impairment is a link between MLL4-inactivated lung tumors and their vulnerability to glycolysis pathway inhibition"

### Supplementary Figure S1

**a**

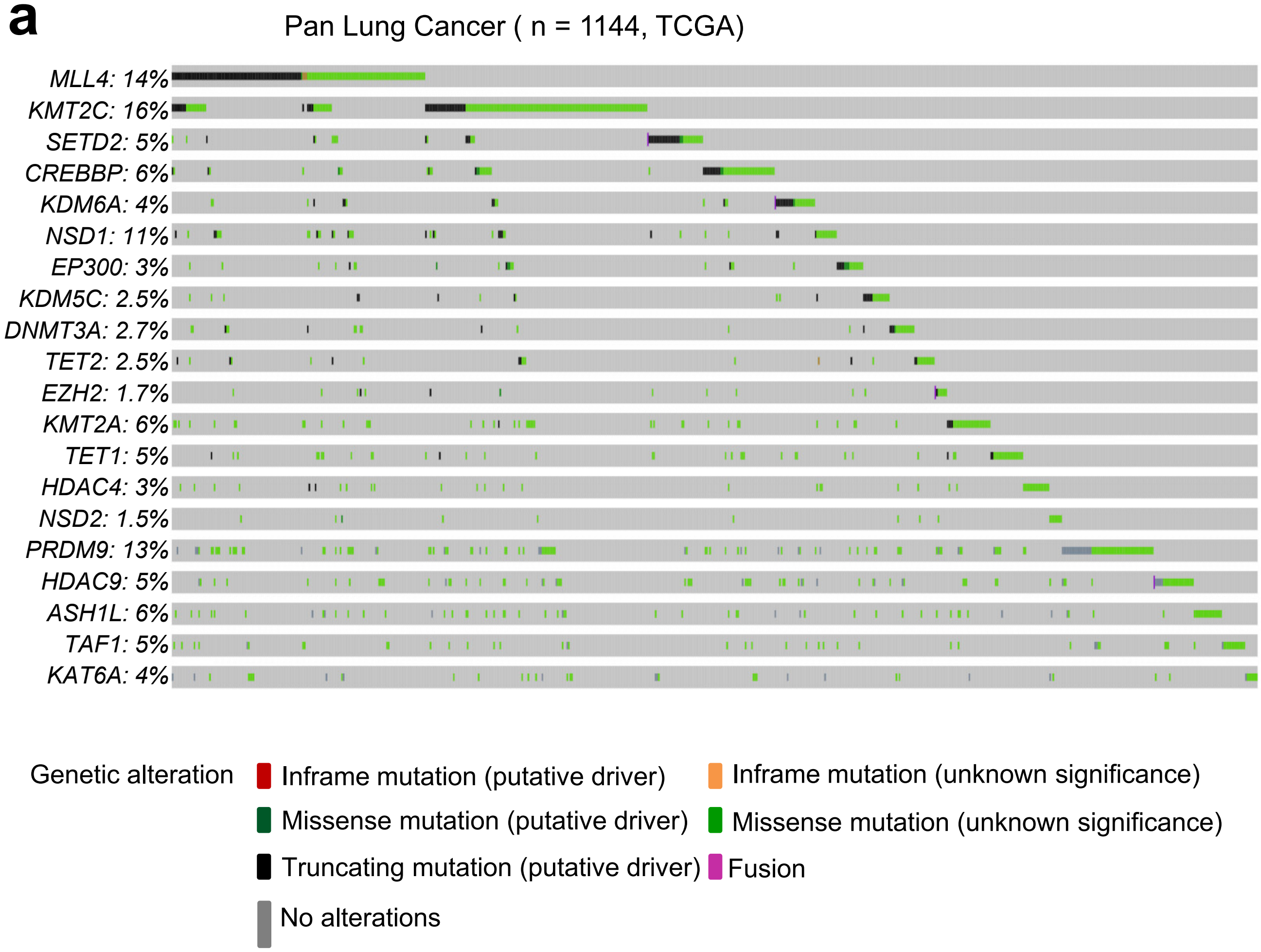


**b**

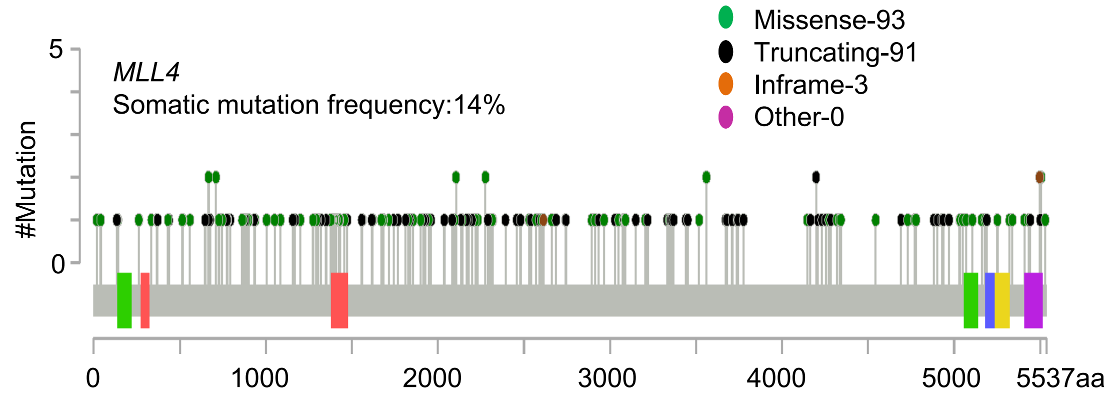


**Supplementary Figure S1**: ***MLL4* is one of the most frequently inactivated genes in human lung cancer samples.** (**a**) The top 20 mutated genes in **Fig. 1a** were analysed using in TCGA Pan-lung cancer (NSCLC) dataset (*n* = 1144) in cBioPortal (http://www.cbioportal.org). Altered samples are mainly shown. (**b**) There was a significantly high percentage (48.7%) of truncating mutations (loss-of-function) in the *MLL4* gene. The lollipop graph shows mutation profiles (missense, truncation, and inframe) in the *MLL4* gene in Pan-lung cancer dataset (*n* = 1144). Data were generated using the TCGA Pan-lung cancer in cBioPortal.

### Supplementary Figure S2

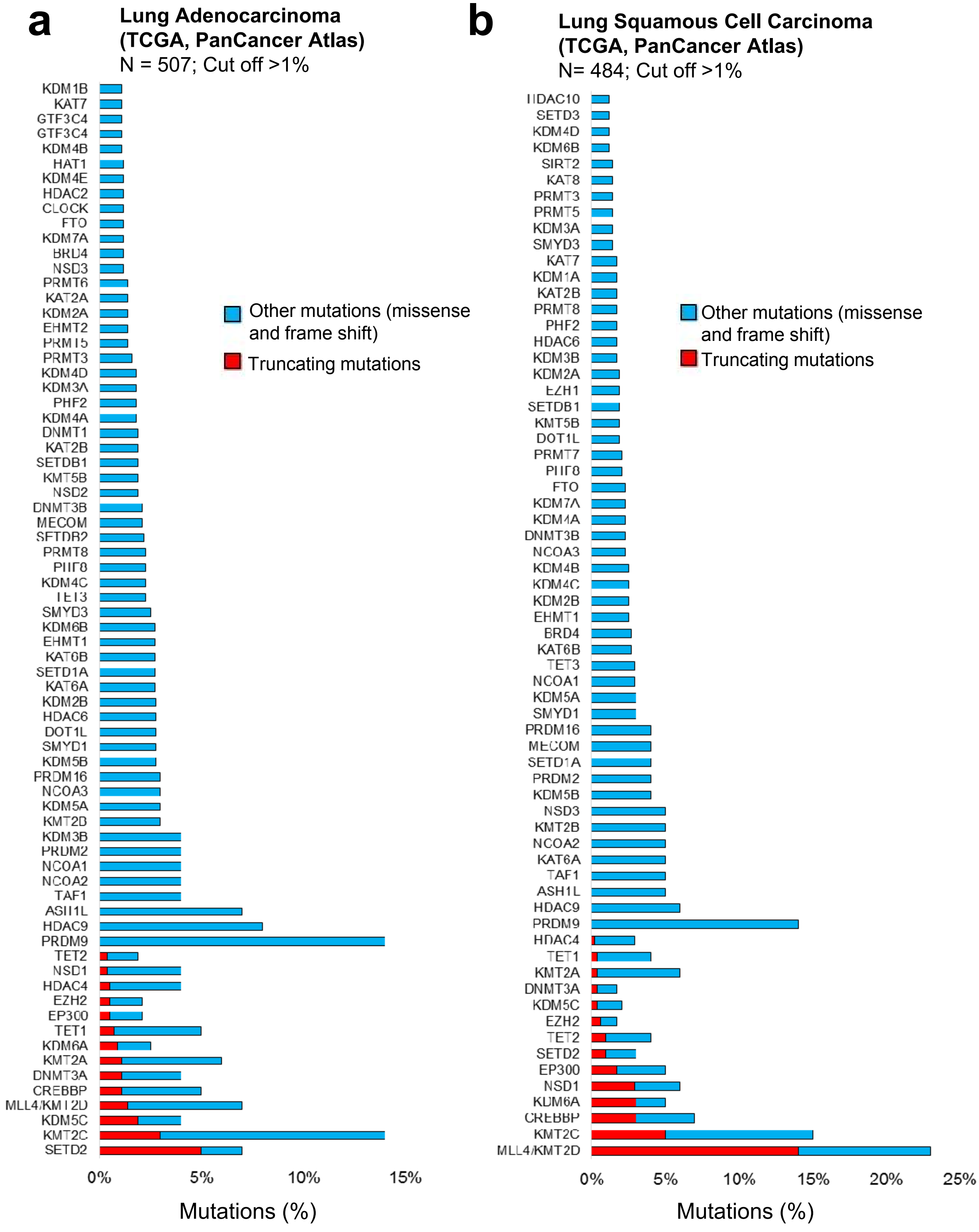


<u>**Supplementary Figure S2**</u>**:** (<u>**a** and **b**</u>) ***MLL4* is one of the most highly inactivated histone methylation modifier in human lung cancer.** Mutations in histone methylation modifiers in the TCGA lung adeno carcinoma (LUAD; n = 507) and lung squamous cell carcinoma (LUSC; 484) dataset were analysed using cBioPortal (http://www.cbioportal.org). Bar graph shows alterations in histone methylation modifiers with more than 1% mutations in LUAD (**a**) and LUSC (**b**) samples. Other mutations represent missense mutations and inframe mutations.

### Supplementary Figure S3

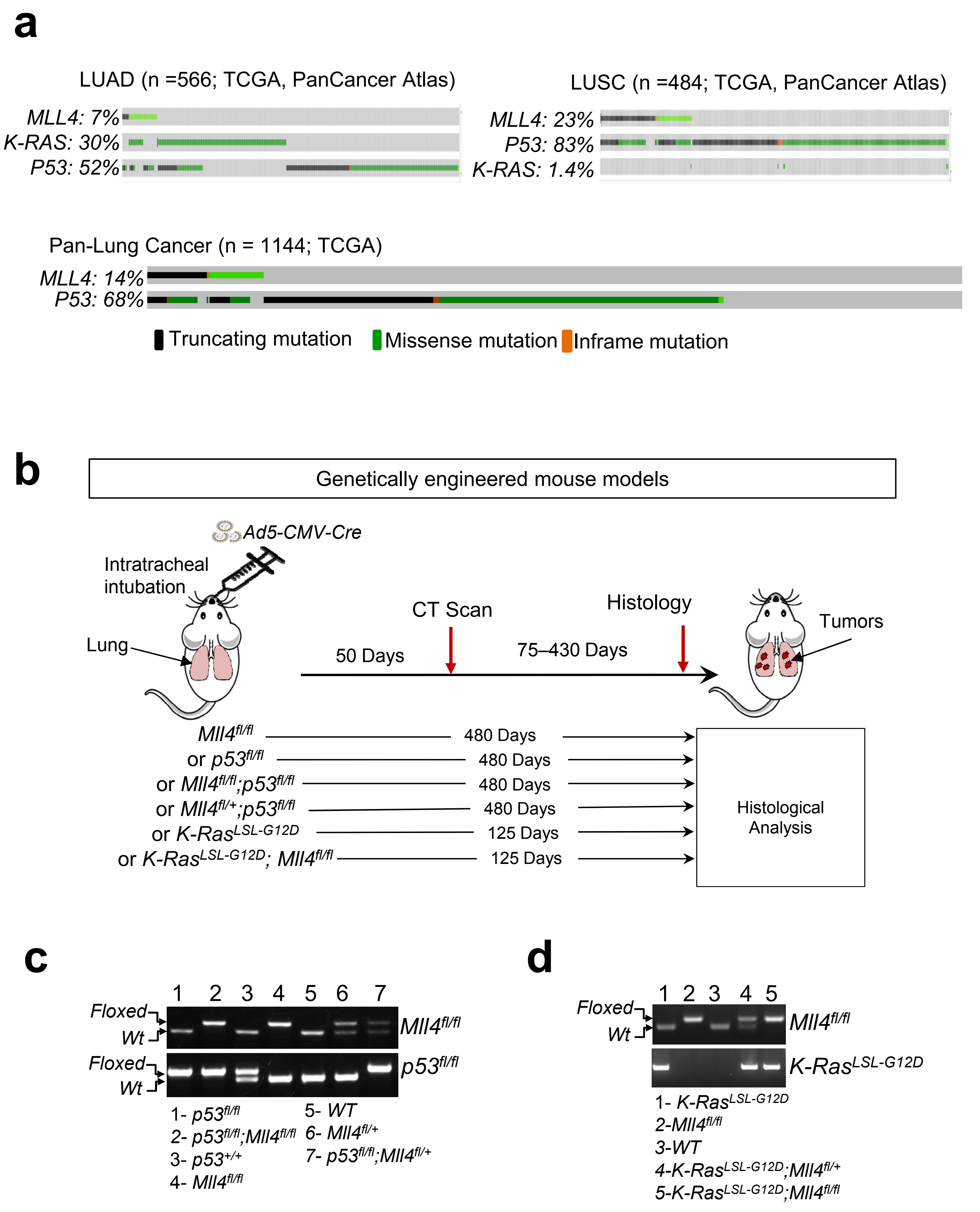


**Supplementary Figure S3**: (**a**) Analysis of the TCGA LUAD and LUSC dataset in cBioPortal showed that *MLL4* mutations often co-occurred with *K-RAS* mutations in human LUAD samples and *p53* mutations in human LUAD and LUSC samples. Altered samples in the TCGA dataset ($n$ = 1144) are mainly shown. (**b**) Our strategy to induce and monitor lung tumorigenesis using new genetically engineered mouse models were summarized. (**c** and **d**) Genotyping results using specific primers showed for the generation of $Mll4^{fl/fl};p53^{fl/f}$ (**c**) and $K\text{-}Ras^{LSL\text{-}G12D};Mll4^{fl/fl}$ (**d**) mice.

### Supplementary Figure S4

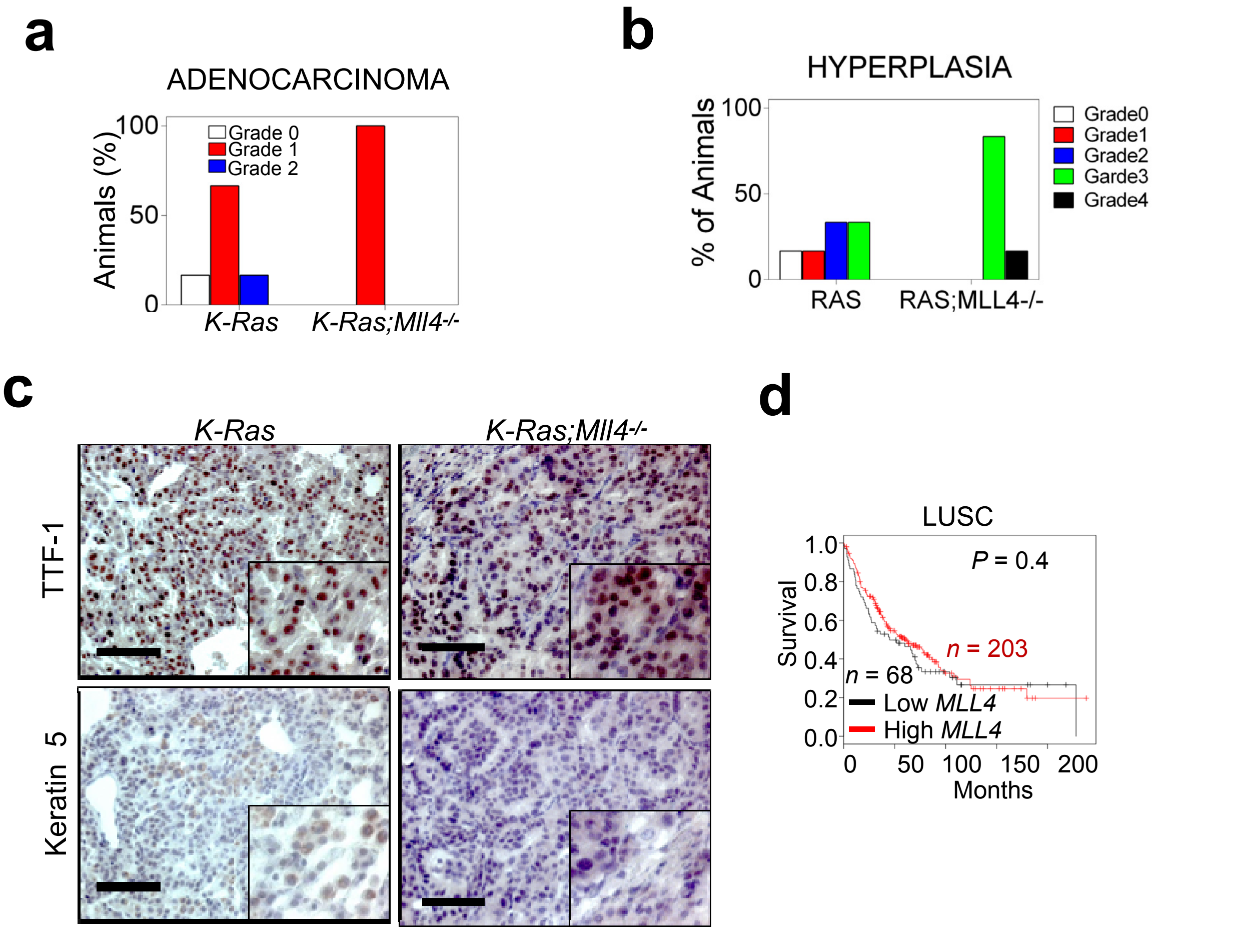


**<u>Supplementary Figure S4</u>: *Mll4* loss accelerates K-Ras-driven LUAD tumorigenesis**. (**<u>a</u>** and **<u>b</u>**) Histopathological analysis of Ad5-CMV-Cre-infected lungs of *K-Ras* and *K-Ras;Mll4*$^{-/-}$ mice showed that *K-Ras;Mll4*$^{-/-}$ mice had a higher percentage of pulmonary parenchyma effaced by lung adenocarcinoma (**a**) and epithelial hyperplasia (**b**) than did those of *K-Ras* mice. Tumors and epithelial hyperplasia grades were based on the percentage of the lung effaced by these lesions; higher grades indicate that a higher percentage of the lung was affected. (**<u>c</u>**) IHC staining of *K-Ras* and *K-Ras;Mll4*$^{-/-}$ lung tumors using TTF-1 and Keratin5, antibodies showed that *K-Ras;Mll4*$^{-/-}$ lung tumors similar to *K-Ras* lung tumors were positive for TTF1 and that *Mll4* loss did not have any obvious effect on low Keratin 5 levels in *K-Ras* lung tumors. TTF1, a lung adenocarcinoma marker; Scale bars, 100 µm. (**<u>d</u>**) The Kaplan-Meier survival analysis showed that low mRNA levels of MLL4 did not correlate with poor survival in LUSC patients. KM Plotter database (http://kmplot.com/analysis) were used for this analysis. Auto cut-off was used to divide low and high groups of samples.

Supplementary Figure S5

**a**

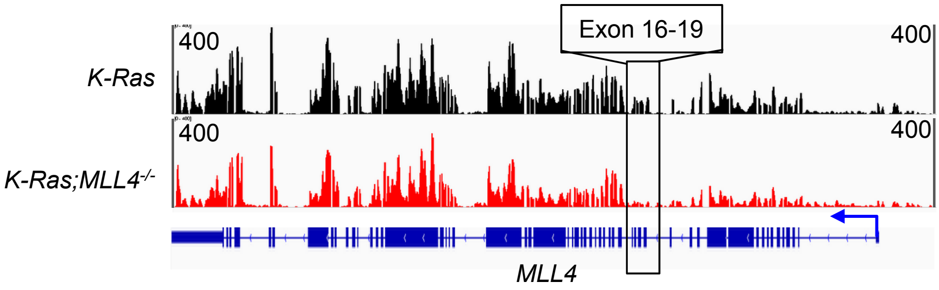


**b**

Pathways downregulated by *Mll4* loss in K-Ras-driven mouse LUAD

| ANNOTATED CELLULAR FUNCTION | SIZE | ES | NES | NOM-p-value | FDR-q-value |
|---|---|---|---|---|---|
| IL6_JAK_STAT3_SIGNALING | 82 | -0.4184 | -1.34824 | 0.039301 | **0.496795** |
| TNFA_SIGNALING_VIA_NFKB | 195 | -0.35153 | -1.25073 | 0.058957 | **0.548042** |
| EPITHELIAL_MESENCHYMAL_TRANSITION | 193 | -0.34841 | -1.23581 | 0.0839 | **0.416064** |
| HEDGEHOG_SIGNALING | 35 | -0.41852 | -1.11486 | 0.309917 | **0.725324** |
| INFLAMMATORY_RESPONSE | 193 | -0.3062 | -1.10957 | 0.202353 | **0.603693** |

**Supplementary Figure S5**: (**a**) Genome browser view of normalized signal of RNA-seq data at *Mll4* locus of *K-Ras* and *K-Ras;Mll4*$^{-/-}$ lung tumor samples. Combined data of two biological replicates from each group are shown. *K-Ras;Mll4*$^{-/-}$ mice showed significant loss of mRNA peaks at exon 16-19. (**b**) Gene set enrichment analysis (GSEA) plot showed that no pathway was significantly downregulated in *K-Ras;Mll4*$^{-/-}$ compared with *K-Ras* tumors..

### Supplementary Figure S6

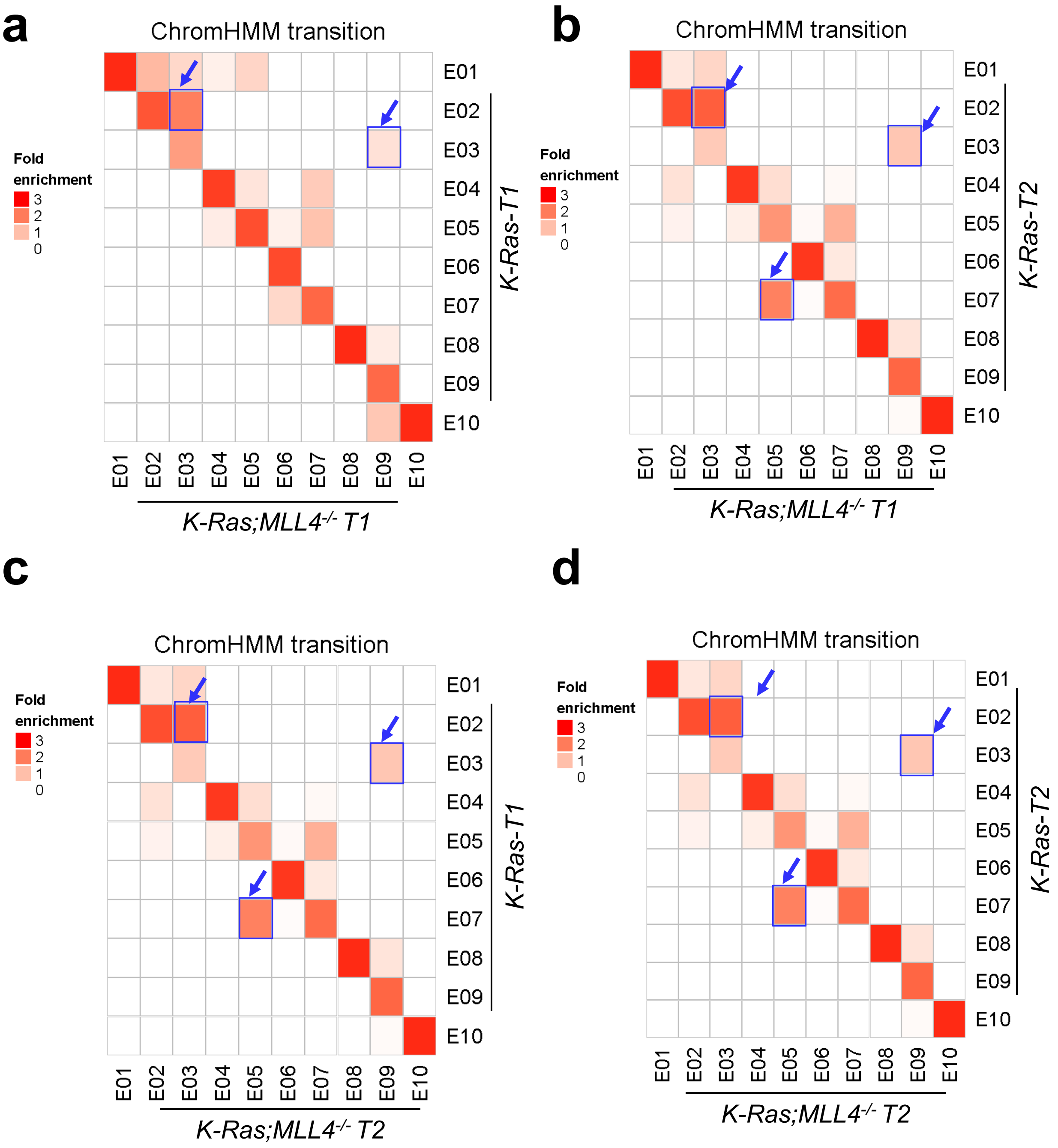


**<u>Supplementary Figure S6</u>: *Mll4* loss weakens active enhancer states in K-Ras–induced lung adenocarcinoma.** (**<u>a</u>**–**<u>d</u>**) chromHMM state transition of the 10-state ChromHMM model was calculated on the basis of six histone modification profiles between *K-Ras;Mll4*$^{-/-}$ and *K-Ras* lung tumors. Heat maps show fold enrichment of transitions of chromatin states between *K-Ras* and *K-Ras;Mll4*$^{-/-}$ lung tumors. The analysis was performed using two different biological replicates. The *K-Ras;Mll4*$^{-/-}$ lung tumors compared with *K-Ras* tumors showed three major transitions in chromatin states from *K-Ras* to *K-Ras;Mll4*$^{-/-}$ lung tumors : 1) E02 (active enhancer) to E03 (weak active enhancer); 2) E03 (weak active enhancer) to E09 (low state); and 3) E07 (transcribed enhancer) to E05 (H3K4me1-low enhancer). T1, tumor 1; T2, tumor 2.

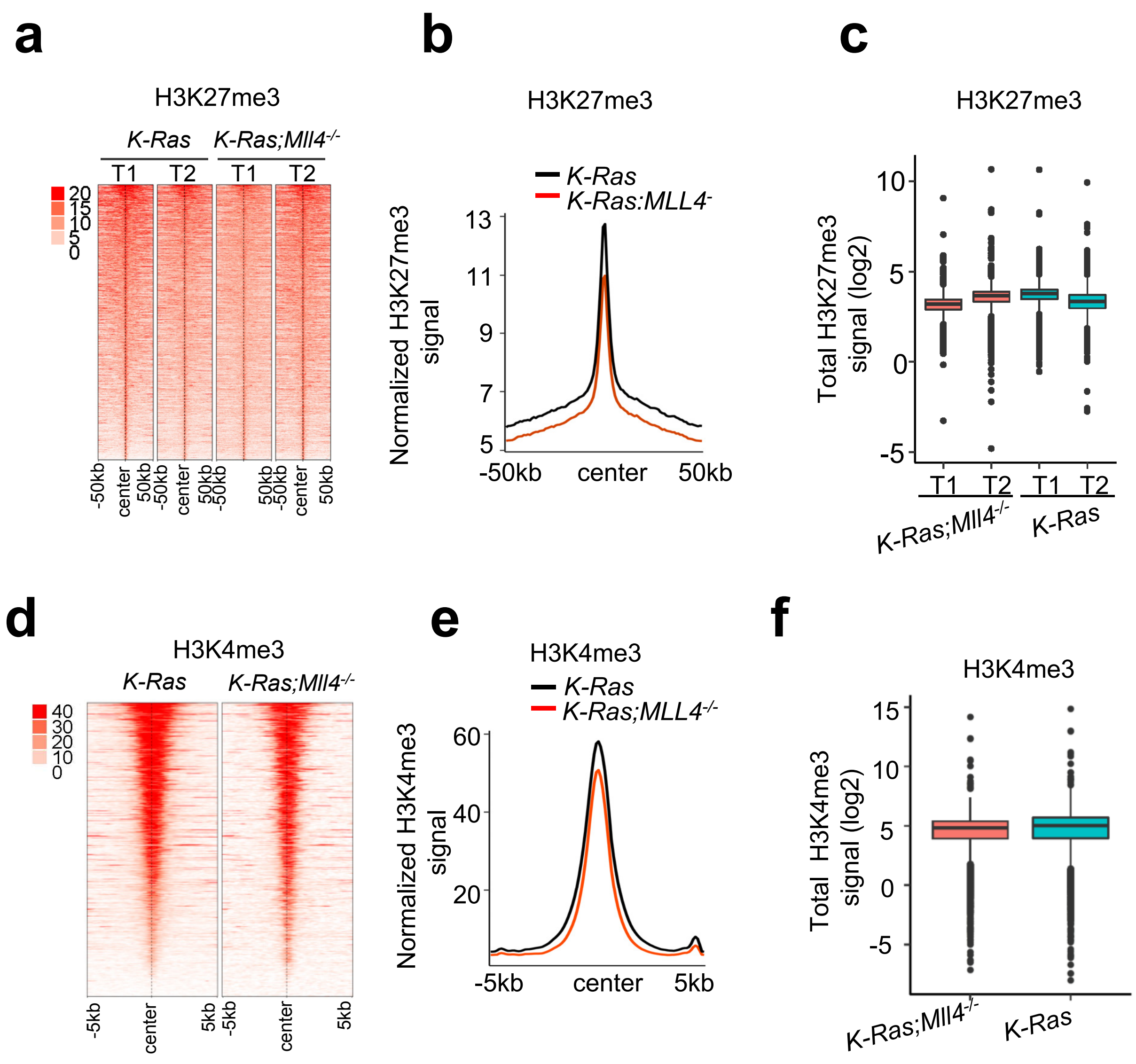


**<u>Supplementary Figure S7</u>: *Mll4* loss does not have an obvious effect on H3K27me3 and H3K4me3 levels in K-Ras–induced lung adenocarcinoma.** (**<u>a</u>**–**<u>c</u>**) There was no significant change in global H3K27me3 levels in Stem Cell Reports. Heat maps (**a**) and average intensity curves (**b**) of ChIP-Seq reads (RPKM) for H3K27me3 were analyzed in a 50kb window centered on the middle of the H3K27me3 peaks in *K-Ras* and *K-Ras;Mll4-/-* lung tumors. H3K27me3 signals ($log_2$RPKM) between *K-Ras* and *K-Ras;Mll4-/-* lung tumors were compared in a box plot (**c**). (**<u>d</u>**–**<u>f</u>**) There was no significant change in global H3K4me3 levels in *K-Ras* and *K-Ras;Mll4-/-* lung tumors. Heat maps (**d**) and average intensity curves (**e**) of ChIP-Seq reads (RPKM) for H3K4me3 were analyzed in a 5kb window centered on the middle of H3K4me3 peaks in *K-Ras* and *K-Ras;Mll4-/-* lung tumors. H3K4me3 signals ($log_2$RPKM) between *K-Ras* and *K-Ras;Mll4-/-* lung tumors were compared in a box plot (**f**).

**a**

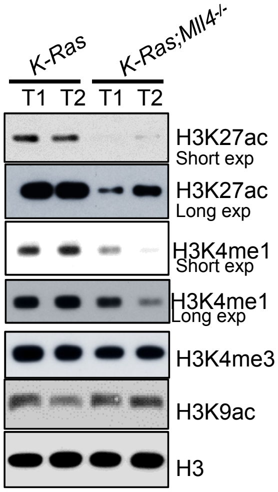


**b**

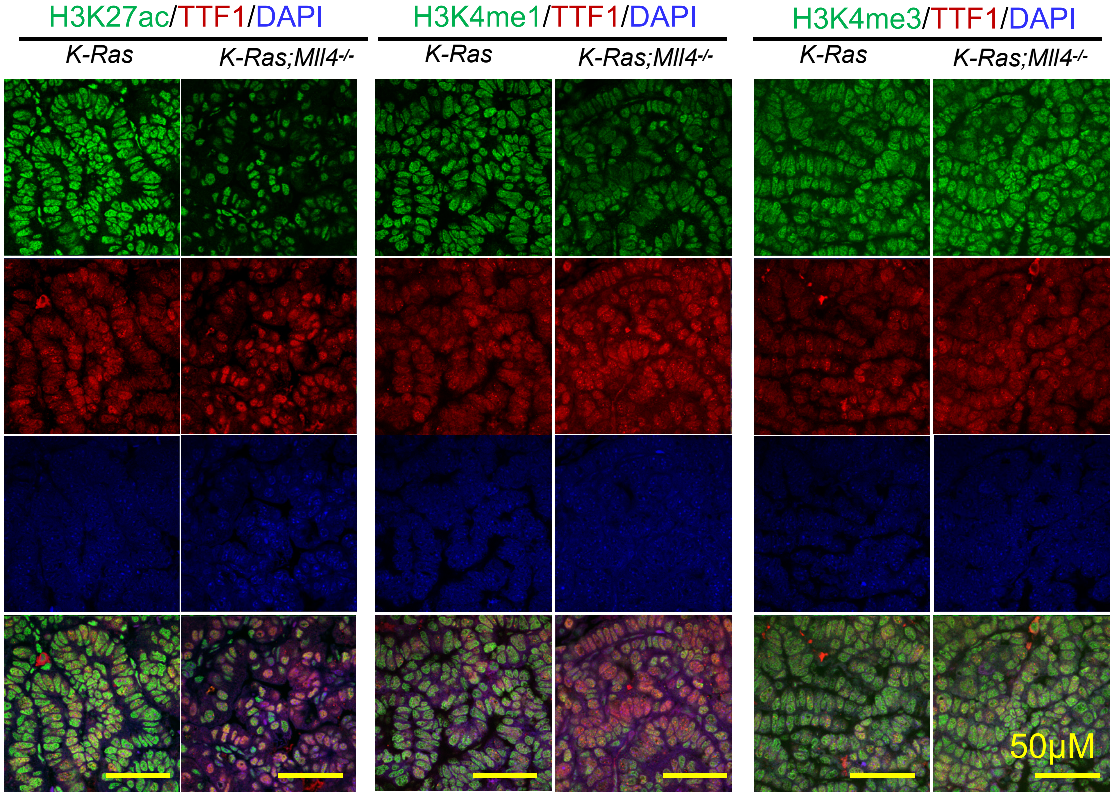


**Supplementary Figure S8**: Western blot analysis (**a**) and immunofluorescence staining (**b**) showed that *Mll4* loss downregulated enhancer signals (H3K27ac and H3K4me1) in K-RAS-induced mouse lung adenocarcinoma. *K-Ras* and *K-Ras;Mll4*$^{-/-}$ mouse lung tumor tissues were analyzed. Yellow scale bars, 50 μm.

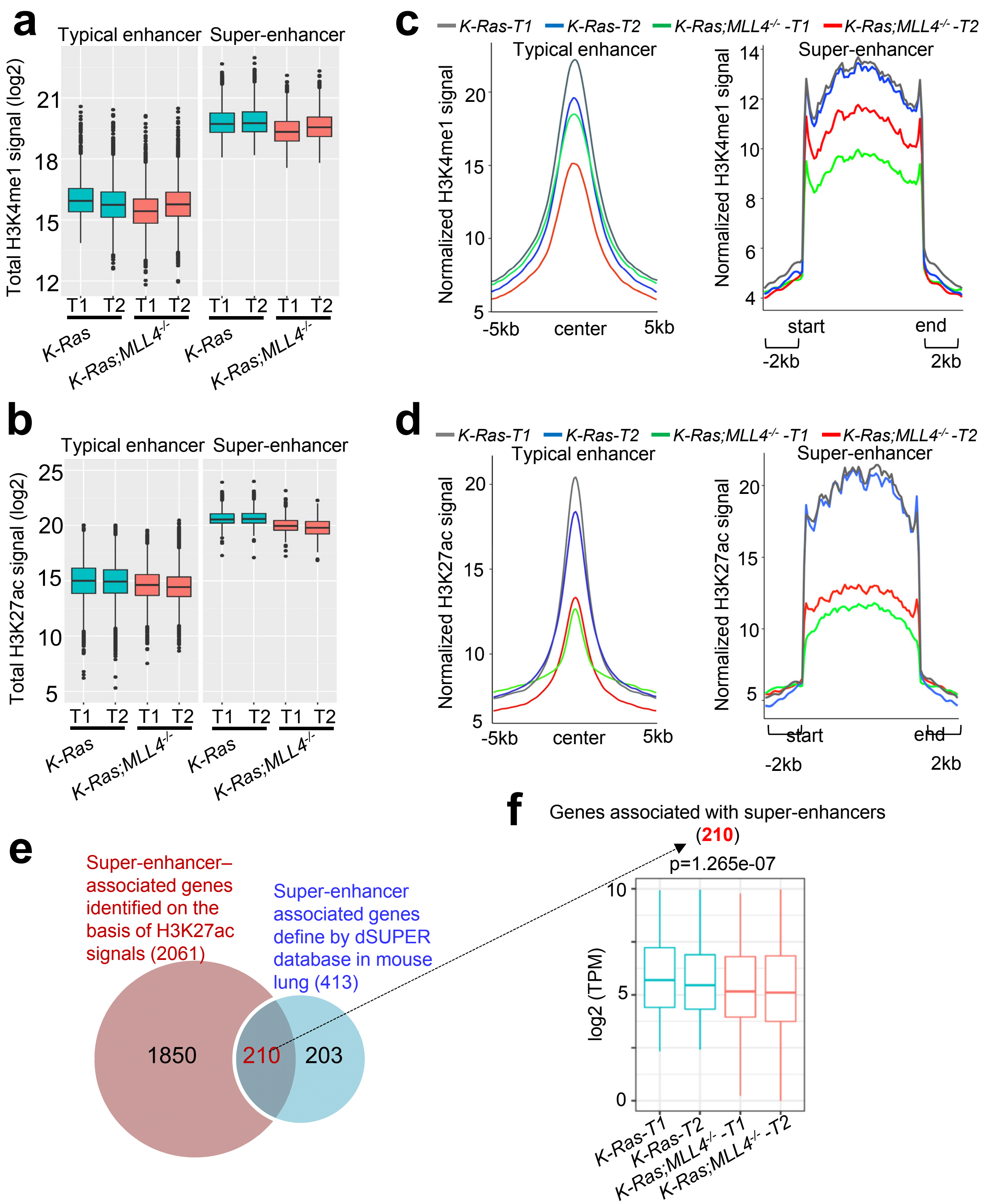


**Supplementary Figure S9**: ***Mll4* loss diminishes more super-enhancer signals than typical enhancer signals and downregulates expression of lung-enriched, super-enhancer–associated genes in K-Ras–induced lung adenocarcinoma.** (**a** and **c**) Analysis of enhancers on the basis of H3K4me1 signals showed that *Mll4* loss diminishes more super-enhancers than typical enhancers. Shown are box plots of H3K4me1 signals for typical enhancers and super-enhancers in *K-Ras* and *K-Ras;Mll4$^{-/-}$* lung tumors (**a**). Average intensities of ChIP-Seq reads for H3K4me1 at the typical enhancer (left panel) and the super-enhancer (right panel) regions were compared between *K-Ras* and *K-Ras;Mll4$^{-/-}$* lung tumors (**c**). (**b** and **d**) Analysis of enhancers on the basis of H3K27ac signals showed that *Mll4* loss diminishes more super-enhancers than typical enhancers. Shown are box plots of H3K27ac signals for typical enhancers and super-enhancers in *K-Ras* and *K-Ras;Mll4$^{-/-}$* lung tumors (**b**). Average intensities of ChIP-Seq reads for H3K27ac at the typical enhancer (left panel) and super-enhancer (right panel) regions were compared between *K-Ras* and *K-Ras;Mll4$^{-/-}$* lung tumors (**d**). (**e**) Venn diagram shows that super-enhancer–associated genes identified on the basis of H3K27ac signals in K-Ras lung tumors substantially overlap with mouse lung super-enhancer genes defined by dbSUPER database (http://bioinfo.au.tsinghua.edu.cn/dbsuper/). (**f**) *Mll4* loss reduced expression of 210 super-enhancer–associated genes in *K-Ras* lung tumors.

### Supplementary Figure S10

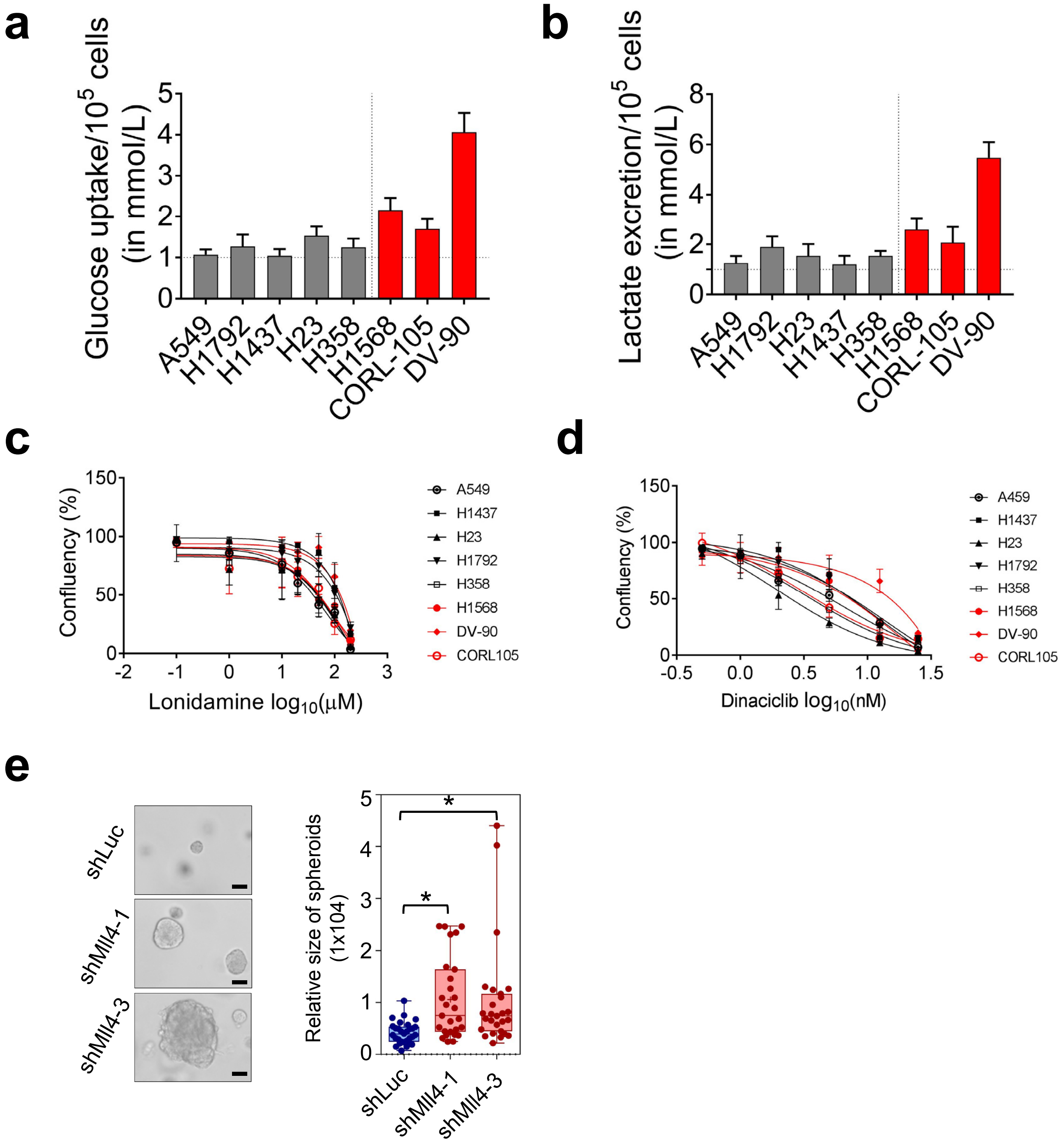


**Supplementary Figure S10**: (**a** and **b**) There were an increased trend of glucose uptake (**a**) and lactate excretion (**b**) in the *MLL4*-mutant cell lines (H1568, DV-90 and CORL105) compared with the *MLL4*-normal cell lines (A459, H1792, H1437, H23, and H358). (**c** and **d**) Cell proliferation inhibition curves showed that Linodamine (**c**) or Dinaciclib (**d**) did not selectively inhibit the proliferation of *MLL4*-normal human lung cancer cell lines (A459, H1437, H23, H1792, and H358) over *MLL4*-mutant human lung cancer cell lines (H1568, DV-90 and CORL105). (**e**) MLL4 knockdown using shmMll4-1 and shmMll4-3 increased spheroid sizes of LKR-10 cells in 3D-culture. Representative images are shown (**left panel**). The boxplot presents the relative size of spheroids formed by MLL4 knockdown (shmMll4-1 and shmMll4-3) LKR-10 cells in 3D-culture (**right panel**). shLuc cells used as a control.

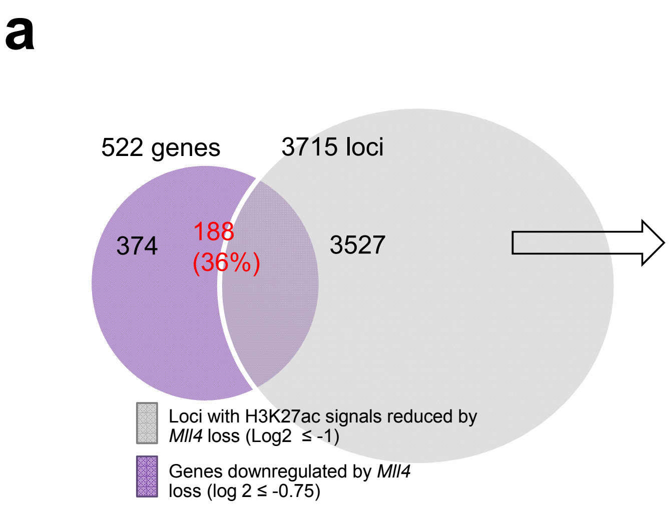


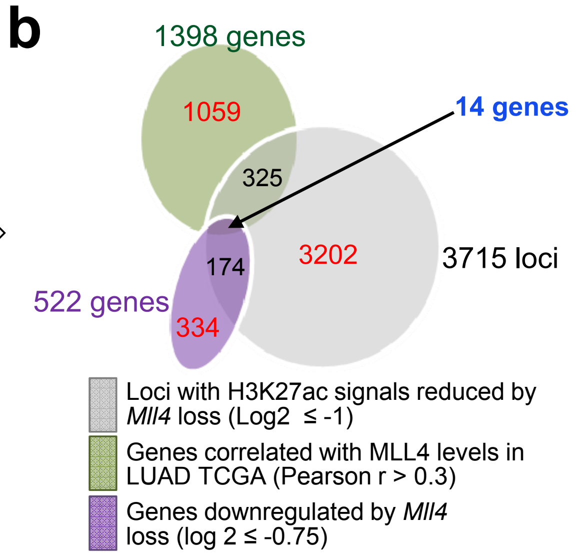


**c** Characteristics of the top 14 genes

| No | GENE | **$Log_2$ FC (RNAseq)** | p-value (RNAseq) | **$Log_2$ FC (H3K27Ac)** | P value (H3K27Ac) | **r (pearson correlation)** | **FC_LUAD (Tumor/ Normal)** | P-VALUE (LUAD) | **SURVIVAL** |
|---|---|---|---|---|---|---|---|---|---|
| 1 | SHANK2 | -2.0252 | 0.005409 | -4.28972 | 4.36E-05 | 0.49 | 0.596 | down*** | Low;Poor |
| 2 | MLANA | -1.74409 | 0.023769 | -1.52758 | 0.414633 | 0.33 | 0.626 | down*** | High;Poor |
| 3 | TRAF3IP1 | -1.54151 | 0.001934 | -1.73023 | 0.07758 | 0.42 | 0.968 | Not sig | Low;Poor |
| 4 | ACACB | -1.16118 | 7.75E-05 | -2.72969 | 0.003311 | 0.53 | 0.312 | down*** | Low;Poor |
| 5 | CRACR2A | -1.06423 | 3.95E-04 | -2.89113 | 0.002267 | 0.3 | 2.03 | Up*** | High;Poor |
| 6 | KIFC3 | -1.04796 | 9.53E-05 | -1.43475 | 0.062539 | 0.33 | 1.03 | Not sig | High;Poor |
| 7 | CASZ1 | -0.96877 | 0.002018 | -1.06683 | 0.156928 | 0.37 | 0.459 | down*** | High;Poor |
| 8 | PER2 | -0.92642 | 0.084606 | -1.66447 | 0.063166 | 0.35 | 0.77 | down*** | Low;Poor |
| 9 | NFASC | -0.89593 | 0.141034 | -2.22851 | 0.033627 | 0.3 | 0.27 | down*** | Low;Poor |
| 10 | CLCN6 | -0.86943 | 5.63E-04 | -2.27149 | 0.016607 | 0.48 | 0.675 | down*** | Low;Poor |
| 11 | TNS2 | -0.852 | 0.014909 | -1.09323 | 0.129067 | 0.31 | 0.326 | down** | Not sig |
| 12 | ankrd23 | -0.81188 | 0.036254 | -1.24506 | 0.252309 | 0.33 | 3.11 | high | Low;Poor |
| 13 | TMEM2 | -0.79389 | 0.046892 | -1.10537 | 0.165816 | 0.33 | 0.66 | down*** | High;Poor |
| 14 | NAV2 | -0.75495 | 0.049805 | -1.82735 | 0.0531 | 0.43 | 0.983 | Not sig | Low;Poor |

**Supplementary Figure S11:** (**a**) A Venn diagram showing the overlapping genes between genes downregulated by *Mll4* loss (n = 522) and genes with H3K27ac ChIP-seq signals reduced by *Mll4* loss (n=3715). (**b**) A Venn diagram showing the overlapping genes between genes downregulated by *Mll4* loss (n = 522), genes with H3K27ac ChIP-seq signals reduced by *Mll4* loss (n=3715), and , genes correlated with *MLL4* expression (n =1398 with r ≥ 0.3) in NSCLC samples (n = 357) in TCGA database. (**c**) Five different characteristics of the top fourteen genes in **Fig. S11b** were analyzed. FC, fold change.

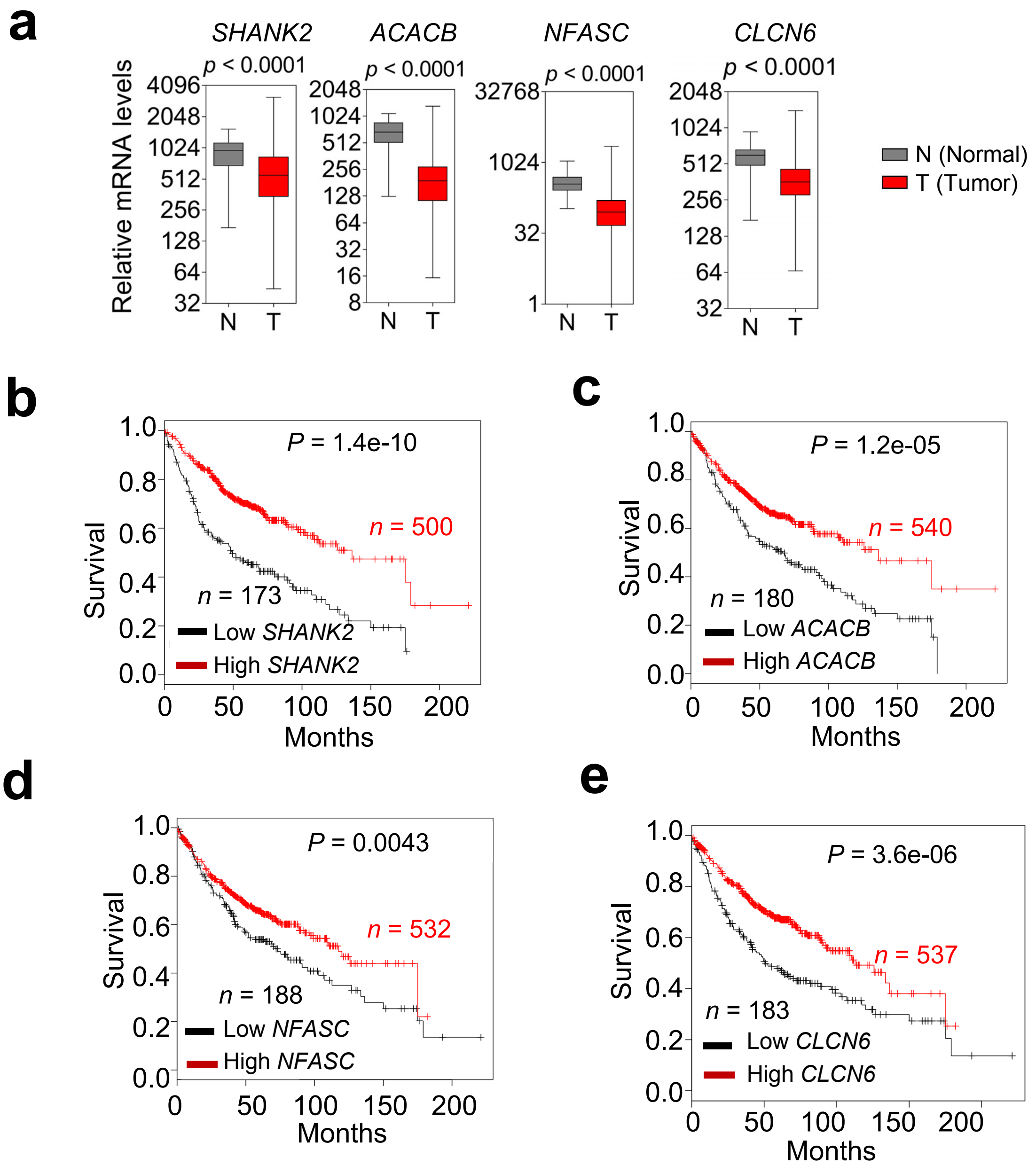


**Supplementary Figure S12**: (**a**) *SHANK2*, *ACACB*, *NFASC*, and *CLCN6* mRNA levels were downregulated in lung adenocarcinoma tumor samples (*n* = 357) compared with their adjacent normal samples (*n* = 54) in TCGA dataset. (**b–e**) Kaplan-Meier survival analysis using the KM Plotter database (http://kmplot.com/analysis) showed that low *SHANK2, ACACB, NFASC* , and *CLCN6* mRNA levels significantly correlated with poor survival of human lung cancer patients. The lower quartile cutoff was used to divide samples into low and high groups. *SHANK2,* probe set 243681_at; *ACACB,* probe set 49452_at; *NFASC,* probe set 213438_at; *CLCN6*, probe set 203950_at.

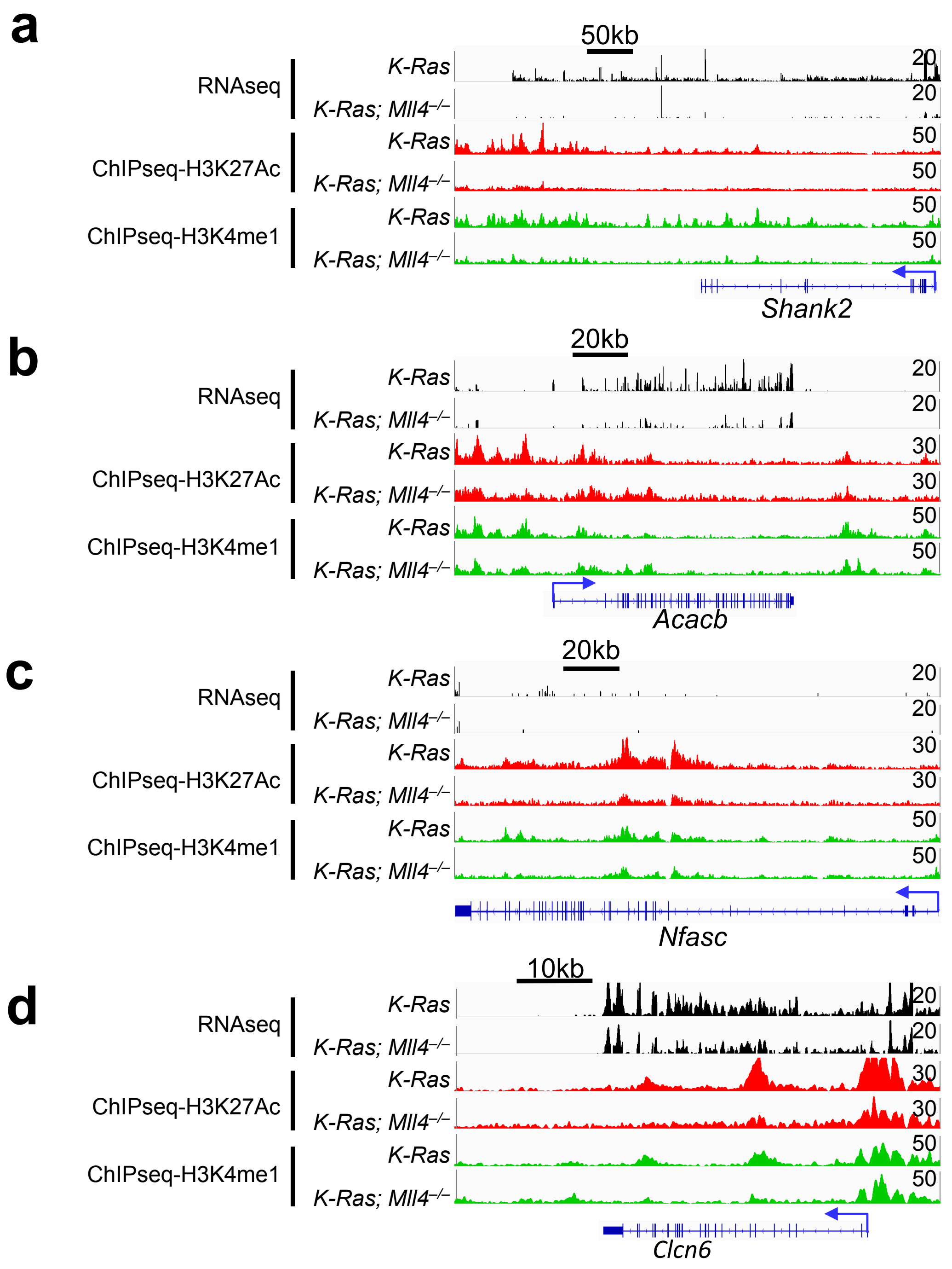


**Supplementary Figure S13**: ***Mll4*** **loss impairs enhancer signals (H3K27ac and H3K4me1) while downregulating gene expression in K-Ras-driven mouse lung tumors**. Genome browser view of normalized signals of RNA-seq data, H3K27ac, and H3K4me1 at *Shank2* (**a**), *Acacb* (**b**), *Nfasc* (**c**), and *Clcn6* (**d**) loci in *K-Ras* and *K-Ras;Mll4$^{-/-}$* lung tumors are shown. All the tracks are average of two biological replicates. H3K27ac and H3K4me1 ChIP-seq signals were normalized to their inputs.

### Supplementary Table S1

Supplementary table S1: Histopathological analysis of Ad5-Cre-infected lung of *Mll4*$^{fl/fl}$ , *p53*$^{fl/fl}$ , *Mll4*$^{fl/+}$;*p53*$^{fl/fl}$, and *Mll4*$^{fl/fl}$;*p53*$^{fl/fl}$ mice.

| **Groups** | ***Mll4*$^{fl/fl}$** | ***p53*$^{fl/fl}$** | ***Mll4*$^{fl/+}$;*p53*$^{fl/fl}$** | ***Mll4*$^{fl/fl}$;*p53*$^{fl/fl}$** |
|---|---|---|---|---|
| Pulmonary adenocarcinomas | 0/5 (0%) | 0/5 (0%) | 0/5 (0%) | 0/5 (0%) |
| Pulmonary adenomas | 0/5 (0%) | 0/5 (0%) | 0/5 (0%) | 0/5 (0%) |
| Multifocal, bronchioloalveolar hyperplasia | 0/5 (0%) | 0/5 (0%) | 0/5 (0%) | 0/5 (0%) |
| BALT hyperplasia | 4/5 (80%) | 4/5 (80%) | 4/5 (80%) | 4/5 (80%) |

### Supplementary Table S2

Supplementary table S2: Mutation status of the *MLL4* gene in human lung cancer cell lines

| Cell line | Chr | Start Position | End Position | Variant Classification | Variant Type | Reference Allele | Tumor Seq Allele | Protein Change |
|---|---|---|---|---|---|---|---|---|
| NCIH1568 | 12 | 49445194 | 49445194 | Nonsense mutation | SNP | C | A | p.E758* |
| DV90 | 12 | 49435199 | 49435199 | Frame_Shift_Del | DEL | G | - | p.Pro2118ProfsTer25(p.P2118fs) |
| CORL105 | 12 | 49434991 | 49434991 | Frame_Shift_Del | DEL | G | - | p.Arg2188ProfsTer74(p.R2188fs) |
| H1437 | 12 | na | na | na | | | | na |
| H1792 | 12 | na | na | na | | | | na |
| H23 | 12 | na | na | na | | | | na |
| H358 | 12 | na | na | na | | | | na |

### Supplementary Table S3

| Supplementary table S1: Primer sequences, shRNA clones and antibodies | | | |
|---|---|---|---|
| | Gene/Protein | | Sequence/clone/company |
| Primers for quantitative RT-PCR (mouse) | Mll4 | Forward | GGC GTT GTG TGG AGT GTA TC |
| | | Reverse | CAC AGT CAT CAC AGA GCA GC |
| | Eno1 | Forward | CAT GGG GAA GGG TGT CTC AC |
| | | Reverse | GTG CCG TCC ATC TCG ATC AT |
| | Pgk1 | Forward | ATG TCG CTT TCC AAC AAG CTG |
| | | Reverse | GCT CCA TTG TCC AAG CAG AAT |
| | Pgam1 | Forward | TCT GTG CAG AAG AGA GCA ATC C |
| | | Reverse | CTG TCA GAC CGC CAT AGT GT |
| | Gapdh | Forward | CATGGCCTTCCGTGTTCCTA |
| | | Reverse | GCCTGCTTCACCACCTTCTT |
| | Ldha | Forward | ACC TCG GTA TTA TTT TTC CAT TTC A |
| | | Reverse | TGT AAT CTT GTT CTG GGG AGC C |
| | Per2 | Forward | GAA AGC TGT CAC CAC CAT AGA A |
| | | Reverse | AAC TCG CAC TTC CTT TTC AGG |
| | Cdk1 | Forward | AGA AGG TAC TTA CGG TGT GGT |
| | | Reverse | GAG AGA TTT CCC GAA TTG CAG T |
| | Ndufa4 | Forward | CGG CTT AGC GTG TGT CCT AA |
| | | Reverse | GCC AAG CGC ATC ACA TAC AG |
| | Ndufa5 | Forward | GAT TGA GCG GGC TTG GGA AA |
| | | Reverse | AAC ATC TGG CTC CTC GTG TG |
| | Ndufa7 | Forward | CCGCTACTCGCGTTATCCAA |
| | | Reverse | TTGGACAGCTTGTGACTGGG |
| | Pdha | Forward | GCA AAC TTG AAG CCA GCC ATC |
| | | Reverse | TCC ACA CCT CTA CAC AGA GC |
| | Gpi1 | Forward | TGG CAA ATC CAT CAC GGA CA |
| | | Reverse | GGA AGT CTC AGG GGA CAA GC |
| | Actin | Forward | GGC TGT ATT CCC CTC CAT CG |
| | | Reverse | CCA GTT GGT AAC AAT GCC ATG T |
| | 18S | Forward | TAGAGGGACAAGTGGCGTTC |
| | | Reverse | CGCTGAGCCAGTCAGTGT |
| Primers for quantitative RT-PCR for Enhancer RNA | E1 | Forward | GTG GGT CCA ACC TCT CCA AG |
| | | Reverse | ATG CTC GCC ATC CAC AAG AA |
| | E2 | Forward | CAA CTG TTT GCC TCT TGC CC |
| | | Reverse | GAG CTG GCT TCC CTT CTC AG |
| Primers for Genotyping | Mll4 flox/flox | Forward (5516_31) | AGAATGGACACTGGAGCTCC |
| | | Reverse (5516_32) | AGAAATCCCCAACCACAGC |
| | p53 flox/flox | Forward (oIMR8543) | GGT TAA ACC CAG CTT GAC CA |
| | | Forward (oimr8544) | GGA GGC AGA GAC AGT TGG AG |
| | Kras-LSL-G12D | Forward(Y117) | CTA GCC ACC ATG GCT TGA GT |
| | | Reverse (Y116) | TCC GAA TTC AGT GAC TAC AGA TG |
| shRNAs Mll4(Sigma) | mouse Mll4 | shmMll4-01 | TRCN0000239234 |
| | | shmMll4-02 | TRCN0000239232 |
| | | shmMll4-03 | TRCN0000239233 |
| | | shmMll4-04 | TRCN0000239231 |
| shRNAs Per2(Sigma) | mouse Per2 | shmPer2-01 | TRCN0000284505 |
| | | shmPer2-02 | TRCN0000096663 |
| | | shmPer2-03 | TRCN0000281782 |
| | | shmPer2-04 | TRCN0000271830 |
| Antibodies (for western, IHC and ChIP) | H3K27ac | | Abcam (ab4729) |
| | H3K27me3 | | Abcam (ab6002) |
| | H3K4me1 | | Cell signaling (5326S) |
| | H3K79me2 | | Abcam (ab3594) |
| | H3K4me3 | | Abcam (ab8580) |
| | H3K9me3 | | Abcam (ab8898) |
| | H3 | | Abcam (ab1791) |
| | H3K16ac | | Abcam (ab4441) |
| | TTF-1 | | Seven Hills Bioreagents (WRAB-1231) |
| | Ki67 | | Cell signaling (9027) |
| | ENO1 | | Abcam (ab227978) |
| | PGK1 | | Abcam (38007) |
| | PGAM1 | | Novus Biological (NBPI-49532) |
| | TTF-1 | | Abcam (ab72876) |
| | MLL4 | | Santa Cruz Biotechnology (sc-2055) |
| | Actin | | Sigma (A5441) |
| | HRP-conjugated anti-mouse-IgG | | Santa Cruz Biotechnology (sc-2055) |
| | HRP-conjugated anti-rabbit-IgG | | Santa Cruz Biotechnology (sc-2004) |
| Antibodies (for IF) | Alexa 488-conjugated anti-mouse IgG |  | Life technologies (A11029) |
|  | Alexa 488-conjugated anti-rabbit IgG |  | Life technologies (A11037) |

**Methods**

**Samples, reagents, and antibodies.** All lung cancer cell lines were procured from ATCC (Rockville, MD, USA), which verifies cell lines using short tandem repeat analysis, and were cultured within 10 – 15 passages. Cell culture reagents and other chemicals were purchased from Gibco, Hyclone, Corning, Sigma-Aldrich and Fisher Bioreagents. The antibodies used for this study are listed in **Supplementary Table S3.**

**Mouse strains and genetically engineered lung cancer models.** The *K-Ras*$^{LSL\text{-}G12D}$ (strain number 01XJ6) and *p53*$^{fl/fl}$ (strain name, B6.129P2-*Trp53*$^{tm1Brn}$/J; stock number, 008462) mice were obtained from the NCI Mouse Repository and Jaxon Laboratory respectively. The *Mll4*$^{fl/fl}$ mice were generated as described earlier [1]. *Mll4*$^{fl/fl}$ mice were crossed with K-*Ras*$^{LSL\text{-}G12D}$ or *p53*$^{fl/fl}$ mice to get desired genotypes for the mouse models of the study. To get *K-Ras*$^{LSL\text{-}G12D}$-*Mll4*$^{fl/fl}$ mice, *Mll4*$^{fl/fl}$ mice were first crossed with K-*Ras*$^{LSL\text{-}G12D}$ mice and the resulting K-*Ras*$^{LSL\text{-}G12D}$;*Mll4*$^{fl/+}$ mice were then crossed with *Mll4*$^{fl/fl}$ To obtain *p53*$^{fl/fl}$;*Mll4*$^{fl/fl}$ mice, *Mll4*$^{fl/fl}$ were first crossed with *p53*$^{fl/fl}$ and the resulting *p53*$^{fl/+}$;*Mll4*$^{fl/+}$ mice were then crossed with *p53*$^{fl/+}$;*Mll4*$^{fl/+}$ to get *p53fl*$^{fl}$;*Mll4*$^{fl/fl}$ mice. The genotype of these mice was confirmed by the regular PCR-based protocol. The primers used for the genotyping are listed in **Supplementary Table S3.**

**In vivo lung tumorigenesis study.** The protocol for induction and monitoring of lung tumorigenesis was used as described previously [2]. In brief, to induce tumor in the mouse lung, 6–8 weeks old mice were infected with 2.5 x $10^7$ Ad5-CMV-Cre virus per mouse by intratracheal intubation method [3]. The tumor progression and survival of mice were compared. For survival analysis, at least 11 mice in each group were used. For ChIP and western blot analysis, the distinct tumors were dissected from the lungs, washed with ice-cold PBS and snap freeze. For histology, IHC, and IF, tumor-bearing lungs were isolated, fixed and processed as previously described [4]. Hematoxylin and eosin (H&E) stained sections of tumor-bearing lungs were evaluated microscopically, and tumors were scored into the different categories 0, I, II, III, and IV on the basis of percentage of pulmonary parenchyma affected by lung adenocarcinoma(s): 0, no tumor present; I, <10% of examined lung affected; II, 10%–20%; III, 21%–50%; IV, >50%. Tumor areas were quantified using ImageScope software.

**Study approval.** The care and use of all mice were approved by the Institutional Animal Care and Use Committee (IACUC) of The University of Texas MD Anderson Cancer Center.

**Micro-CT.** The mice were monitored for tumor growth using micro-CT as previously described [2]. Briefly, the mice were anesthetized with a dose of 5% isoflurane and maintained at 2% isoflurane. The mice were intubated using a 20 gauge x 1-inch catheter and were transferred onto the XRad 225Cx (Precision X-Ray Corporation). The mice were mechanically ventilated in a small animal ventilator, and micro-CT images were captured at 60 KvP, 4 mA, and 3 rpm. Animal's breathing was held at 20cmH20 during the 20-second acquisition. Three to five mice (30 days post-infection) per group were monitored by micro-CT.

**H&E staining, immunohistochemistry (IHC) experiments, and immunofluorescence (IF).** The tumor-bearing lungs were isolated and fixed with 10% formalin buffer. The fixed lung tissues were embedded in paraffin and were cut into 8 μm thick sections. For histological examination, a standard hematoxylin/eosin staining was performed. IHC and IF experiments were performed as described previously [2]. Briefly, sections were subjected to antigen retrieval (antigen retrieval solution, Vector Laboratories, Burlingame, CA) followed by blocking in 10% horse serum for 1 hour at RT. For IF, Alexa 488-conjugated anti-mouse IgG and Alexa 568-conjugated anti-rabbit IgG secondary antibodies were used for detection, and images of tumor regions were captured using a laser confocal microscope. For the quantification of IF staining, signal intensities of glycolytic enzymes in TTF1-positive tumor cells were measured using ImageJ. The primary antibodies used for IHC are listed in **Supplementary Table S3**.

**RNA isolation, quantitative RT-PCR and Western blot analysis of lung tumor cells.** The distinct tumor tissues were dissected and cut into <1mm pieces. The dissected tumor tissues were then digested in Collagenase type 1 and DNAase1 for 45 min followed by 0.25% trypsin for 10 min. To remove red blood cells, the digested single cell suspension was then treated with red blood cell lysis buffer for 2–3 min. To further enrich tumor cells, CD45-positive cells were then removed using MagniSort™ Mouse CD45 Depletion Kit (ThermoFisher Scientific). The depletion of CD45-positive cells was confirmed by flow cytometry analysis. Total RNA was isolated using Trizol reagent (Life Technologies).

Reverse transcription (RT)-PCR and Western blot analysis were performed as described earlier [2]. In brief, for quantitative RT-PCR, iQ SYBR Green Supermix (BioRad) was used for PCR amplification and signals were acquired using CFX384 real-time PCR detection system (BioRad). β-Actin mRNA or 18s rRNA levels were used as internal control. Each experiment was performed in triplicate. The primers and antibodies used for quantitative RT-PCR and ChIP assays are listed in **Supplementary Table S3.**

**RNA-Seq analysis.** The RNA isolated from CD45-depleted tumor tissue samples were sequenced using the Illumina HiSeq 2000. The RNAseq data were processed by pyflow-RNAseq (https://github.com/crazyhottommy/pyflow-RNAseq), a snakemake based RNAseq pipeline. Raw reads were mapped by STAR [5], RPKM normalized bigwigs were generated by deeptools [6], and gene counts were obtained by featureCount [7]. Differentially expression analysis was carried out using DESeq2 [8]. DAVID (version 6.8) was employed for Gene Ontology (GO) analysis as described previously [2]. Gene Set enrichment analysis was done using the GSEA [9] tool from Broad Institute. The pre-rank mode was used. The signed fold change *–log10 (p-value) metric was used for pre-ranking the genes.

**TCGA RNAseq data analysis.** TCGA lung adenocarcinoma (LUAD) and lung squamous carcinoma (LUSC) RNAseq raw counts were downloaded using TCGAbiolinks [10]. The mutation MAF files were downloaded with TCGAbiolinks as well. Mutation status of was inferred from the MAF files. 40 LUSC and LUAD high expressed wild-type samples and 40 MLL4 nonsense tumors (see supplementary data for samples included in the analysis) were compared using DESeq2, the signed fold change *–log10(p-value) metric was used to pre-rank the gene list and for GSEA pre-rank analysis. 20 most highly expressed MLL4 wild-type tumors and 20 most lowly expressed MLL4 wild-type tumors in the LUAD cohort was compared using DESeq2. The signed fold change *–log10 (p-value) metric was used to pre-rank the gene list and for GSEA pre-rank analysis.

**Expression and survival analysis.** The LUAD datasets of TCGA were used for expression and survival analysis. Oncoprint and correlation data were analyzed using the cBio cancer genomics portal (http://www.cbioportal.org) [12-14]. For the survival analysis, a publicly available LUAD transcriptomic datasets were used by http://kmplot.com/analysis/index.php?p=service&cancer=lung website [15].

**ChIP-Seq assays.** Chromatin immunoprecipitation for lung tumor tissue was performed with minor modifications of a previous procedure [16]. Briefly, distinct lung tumor tissues (3 mg per antibody) were dissected from the lungs and cut into 1mm pieces, homogenized using MACS dissociator and cross-linked using 1% paraformaldehyde for 10 min at 37°C. Crosslinking was then stopped by adding 0.125M glycine for 5mins, and tissues were washed with PBS and stored at -80°C. Later, tissues were thawed on ice and lysed with ChIP harvest buffer (12 mM Tris-Cl, 0.1x PBS, 6 mM EDTA, 0.5% SDS) for 10min on ice. Sonication conditions were optimized for lung tumor tissues using bioruptor sonicator to achieve a shear length of 250–500bp. Antibody-dynabead mixtures were incubated for 1 hr at 4°C and tissue extracts were then incubated overnight with antibody-dynabead mixtures. After overnight incubation, immunecomplexes were washed in following order: 5 times with RIPA buffer, twice with RIPA-500 (RIPA with 500mM NaCl) and twice with LiCl wash buffer (10mM Tris-HCl pH8.0, 1mM EDTA pH8.0, 250mM LiCl, 0.5% NP-40, 0.1% DOC). For decrosslinking and elution, immunecomplexes were incubated overnight at 65$^{o}$C in direct elution buffer (10mM Tris-Cl pH8.0, 5mM EDTA, 300mM NaCl, 0.5% SDS). Eluted DNA was then treated with Proteinase K (20mg/ml) and RNaseA and DNA clean-up was done using SPRI beads (Beckman-Coulter). Library was prepared as described earlier [16] using NEB adapters. Libraries were multiplexed together and sequencing was performed in Hiseq2000 or HiSeq4000 (Illumina).

**ChIP-seq analysis.** ChIP-seq data were quality controlled and processed by pyflow-ChIPseq [16], a snakemake based ChIPseq pipeline [17]. Briefly, raw reads were mapped by bowtie1 [18] and duplicated reads were removed. Only uniquely mapped reads were retained. RPKM normalized bigwigs were generated by deeptools [6] and the tracks were visualized with IGV [19]. Peaks were called using MACS1.4 [20] with a p-value of 1E-8. Chromatin state was called using ChromHMM [21] and the emission profile was plotted by ComplexHeatmap [22]. Heatmap was generated by R package EnrichedHeatmap. ChIP-seq peaks were annotated with the nearest genes using ChIPseeker [23]. Super-enhancers were identified using ROSE [24] based on H3K27ac ChIP-seq data.

**ChromHMM transition:** ChromHMM profiles of two *Mll4*$^{-/-}$;*K-Ras* and two *K-Ras* samples are consolidated using epilogos (https://github.com/Altius/epilogos). A pipeline was made to automate the calculation and scripts used to re-code the chromHMM states can be found https://github.com/crazyhottommy/pyflow-chromForest/tree/vsurf_merge. With the output of Epilogos, the chromatin state for each bin was chosen for the state that contained the greatest weights. A helper script can be found in the above link. The output for each group was analyzed by java -mx12000M -jar ChromHMM.jar OverlapEnrichment. The matrix output from OverlapEnrichment was scaled by columns and plotted using ComplexHeatmap (https://bioconductor.org/packages/release/bioc/html/ComplexHeatmap.html).

**Three-dimensional cell culture.** The three-dimensional (3D) cultures were adapted from the procedures previously described [25]. In brief, 96-well plate coated with 6 µl of Engelbreth-Holm-Swarm tumor matrix (Matrigel, BD Biosciences) and kept on RT for 30 min. The cells were trypsinized and suspended in DMEM medium containing 50% Matrigel. 2 x $10^5$ cells were seeded per well on the coated 96-well plates. Every third day, the cells were replenished with fresh DMEM medium containing 10% FBS. The cultures were maintained for 10-14 days, and images were captured.

**Stable knockdown, overexpression and rescue experiments.** For knockdown experiments, lentivirus-based, puromycin-resistant shRNAs were purchased from Sigma (**Supplementary Table S3**). The shRNA-infected cells were selected in puromycin-containing medium (1 µg/ml). shLuciferase (shLuc)-infected cells were used as a control. For ectopic overexpression and rescue experiments, human Per2 cDNA was cloned into the lentivirus vector pLenti6.3/V5-DEST (Thermo Fisher Scientific) using standard cloning methodology. Cells infected with pLenti-Per2 were selected in blasticidin-containing medium (2 µg/ml). pLenti-GFP infected-cells were used as controls.

**Glucose uptake and lactate excretion assay:** Cells were seeded in triplicate 12-well plates. The wells without cells were used as a baseline reading. On the second day, media of each well were changed with 1 ml fresh media (including the wells without cells). After 48 hrs, 600ul media were collected from each well. Media was centrifuged at 3000rpm for 3-5 min at 4C. 200 ul media were transferred into 96 well plate and glucose and lactate levels were measured using YSI according to manufacturer's protocol. Their levels were normalized with the cell number.

**Cell line inhibitor experiments.** Cells were seeded at a density of 1.5x$10^3$ cells in four replicates in 96- well plates. Plated cells were then treated with different concentration of inhibitors. Human LUAD *MLL4* wild-types (A549, H1792, H23, H1437 and H358) and *MLL4* mutants (H1568, DV-

90 and CORL-105) cells were treated with a range of concentrations of 2-DG (1 to 1000 μM), POMHEX (0.05 to 2 μM), SAHA (0.1 to 20 μM) and AR-42 (1 to 500 nM) for one week. The cells were replenished with inhibitor containing medium on every alternate day. After 7 days, the cell growth was quantified using Celigo followed by crystal violet staining. DMSO treated cells were used as a vehicle controls.

**Drug treatment of xenograft mouse models.** Cells ($5 \times 10^6$) in 100 μl of Matrigel were subcutaneously injected in both the flanks of 6 to 8-week-old athymic nu/nu mice. After 8 days when tumors became palpable, mice bearing tumors were randomly separated into two groups. Mice were treated with intraperitoneal injections of 500mg/kg body weight of 2-DG on an alternate day for 20 days. The drug was prepared in sterile water hence sterile water was injected as a vehicle control. Tumors were measured on every alternate day by caliper. Tumor volume was calculated using the ellipsoid volume formula (1/2 × l × w × h) as described earlier [4]. After 20 days of treatment, the mice were euthanized and tumors were collected for histology.

**Statistical analysis.** For correlation analysis, the chi-squared test was performed to calculate the level of significance. The two-sided log-rank method was used to test the statistical significance of survival data using IBM SPSS Statistics 23. The two-tailed Student's t-test was used to determine the statistical significance of two groups of data using GraphPad Prism. Data are presented as means ± standard error of the mean (SEM; error bars) of at least three independent experiments or three biological replicates. *P*-values less than 0.05 were considered statistically significant. *, $P$ <0.05; **, $P$ <0.01; and ***, $P$ <0.001 indicate statistically significant differences.

**Data availability**

RNA-seq and ChIP-seq data that support the findings of this study have been deposited in GEO database with the accession codes GSE116658 (token only for reviewers: qdirksoehrqlpwh) and GSE116659 (token only for reviewers: spynwacojbwbzqd).

**References:**


1 Dhar, S. S. *et al.* MLL4 Is Required to Maintain Broad H3K4me3 Peaks and Super-Enhancers at Tumor Suppressor Genes. *Molecular cell* **70**, 825-841 e826, (2018).

2 Alam, H. *et al.* HP1gamma promotes lung adenocarcinoma by downregulating the transcription-repressive regulators NCOR2 and ZBTB7A. *Cancer Res*, (2018).

3 DuPage, M., Dooley, A. L. & Jacks, T. Conditional mouse lung cancer models using adenoviral or lentiviral delivery of Cre recombinase. *Nat Protoc* **4**, 1064-1072, (2009).

4 Wagner, K. W. *et al.* KDM2A promotes lung tumorigenesis by epigenetically enhancing ERK1/2 signaling. *The Journal of clinical investigation* **123**, 5231-5246, (2013).

5 Dobin, A. *et al.* STAR: ultrafast universal RNA-seq aligner. *Bioinformatics* **29**, 15-21, (2013).

6 Ramirez, F. *et al.* deepTools2: a next generation web server for deep-sequencing data analysis. *Nucleic Acids Res* **44**, W160-165, (2016).

7 Liao, Y., Smyth, G. K. & Shi, W. featureCounts: an efficient general purpose program for assigning sequence reads to genomic features. *Bioinformatics* **30**, 923-930, (2014).

8 Love, M. I., Huber, W. & Anders, S. Moderated estimation of fold change and dispersion for RNA-seq data with DESeq2. *Genome biology* **15**, 550, (2014).

9 Subramanian, A. *et al.* Gene set enrichment analysis: a knowledge-based approach for interpreting genome-wide expression profiles. *Proceedings of the National Academy of Sciences of the United States of America* **102**, 15545-15550, (2005).

10 Colaprico, A. *et al.* TCGAbiolinks: an R/Bioconductor package for integrative analysis of TCGA data. *Nucleic Acids Res* **44**, e71, (2016).

11 Mularoni, L., Sabarinathan, R., Deu-Pons, J., Gonzalez-Perez, A. & Lopez-Bigas, N. OncodriveFML: a general framework to identify coding and non-coding regions with cancer driver mutations. *Genome biology* **17**, 128, (2016).

12 Cerami, E. *et al.* The cBio cancer genomics portal: an open platform for exploring multidimensional cancer genomics data. *Cancer discovery* **2**, 401-404, (2012).

13 Cancer Genome Atlas Research, N. Comprehensive molecular profiling of lung adenocarcinoma. *Nature* **511**, 543-550, (2014).

14 Gao, J. *et al.* Integrative analysis of complex cancer genomics and clinical profiles using the cBioPortal. *Science signaling* **6**, pl1, (2013).

15 Gyorffy, B., Surowiak, P., Budczies, J. & Lanczky, A. Online survival analysis software to assess the prognostic value of biomarkers using transcriptomic data in non-small-cell lung cancer. *PLoS One* **8**, e82241, (2013).

16 Terranova, C. *et al.* An Integrated Platform for Genome-wide Mapping of Chromatin States Using High-throughput ChIP-sequencing in Tumor Tissues. *Journal of visualized experiments : JoVE*, (2018).

17 Koster, J. & Rahmann, S. Snakemake--a scalable bioinformatics workflow engine. *Bioinformatics* **28**, 2520-2522, (2012).

18 Langmead, B., Trapnell, C., Pop, M. & Salzberg, S. L. Ultrafast and memory-efficient alignment of short DNA sequences to the human genome. *Genome biology* **10**, R25, (2009).

19 Robinson, J. T. *et al.* Integrative genomics viewer. *Nature biotechnology* **29**, 24-26, (2011).

20 Zhang, Y. *et al.* Model-based analysis of ChIP-Seq (MACS). *Genome biology* **9**, R137, (2008).

21 Ernst, J. & Kellis, M. ChromHMM: automating chromatin-state discovery and characterization. *Nat Methods* **9**, 215-216, (2012).

22 Gu, Z., Eils, R. & Schlesner, M. Complex heatmaps reveal patterns and correlations in multidimensional genomic data. *Bioinformatics* **32**, 2847-2849, (2016).

23 Yu, G., Wang, L. G. & He, Q. Y. ChIPseeker: an R/Bioconductor package for ChIP peak annotation, comparison and visualization. *Bioinformatics* **31**, 2382-2383, (2015).

24 Loven, J. *et al.* Selective inhibition of tumor oncogenes by disruption of super-enhancers. *Cell* **153**, 320-334, (2013).

25 Lee, G. Y., Kenny, P. A., Lee, E. H. & Bissell, M. J. Three-dimensional culture models of normal and malignant breast epithelial cells. *Nature methods* **4**, 359-365, (2007).
